# *sox17* mutagenesis reveals microglia can be impacted by genetic maternal effects

**DOI:** 10.1101/2025.07.30.667678

**Authors:** Camden A. Hoover, John Dennen, Dailin Gan, Jun Li, Cody J. Smith

## Abstract

Embryonic development is precisely shaped by maternal and zygotic factors. These maternal factors exert their influence through maternal effects, a phenomenon where an offspring’s phenotype is determined, at least in part, by the mother’s environment and genotype. While environmental maternal effects can cause phenotypes that present both early and later in life, genetic maternal effects generally induce phenotypes in the earliest embryonic stages. Here, we reveal a genetic maternal effect that influences the development of cells that arise after early embryogenesis, highlighting that specific cell types can be susceptible to late-onset genetic maternal effects. Using zebrafish to study microglia, the resident immune cells of the brain, we identified a mutation in *sry-related HMG box gene-17 (sox17)* that exhibits a maternal effect phenotype that presents as a reduction of microglia in the brain and precursors in the yolk sac. We demonstrate that *sox17* is expressed in microglia and their yolk sac precursors and is maternally-loaded. We show that *sox17* restoration via embryonic injection reverses the maternal effect on microglia and yolk sac cells in *sox17* mutants. To identify additional genes interacting with *sox17*, we nominated genes from scRNA sequencing analysis of mouse embryonic microglia to perform a genetic screen using CRISPR mutagenesis and a custom-built robot that captures confocal images of the zebrafish brain in high-throughput. This screen identified *f11r.1, gas6,* and *mpp1* as modifiers of microglia abundance in the embryonic brain, which we demonstrated are also expressed in zebrafish microglia. Transcriptional and mutant analyses with these new modifiers suggest that *sox17* positively regulates *mpp1* transcription. These results demonstrate that microglia are susceptible to genetic maternal effects, in addition to their known sensitivity to environmental maternal effects. Our findings reveal a late-onset phenotype associated with the maternal genotype, expanding the recognized impact of genetic maternal effects beyond initial embryo viability and into long-term vigor.

## INTRODUCTION

The construction of a precisely organized embryo is influenced by genetic and maternal effects^1–7^. Maternal effects are phenotypes present in progeny that are determined, at least in part, by the mother’s genotype^2,3,8^. Maternal effects can be induced by both genetic and environmental factors^3,4,6,8^. The genes that influence maternal effect phenotypes in offspring are called maternal effect genes. Maternal effect genes are typically involved in biological processes prior to zygotic genome activation and their disruption often induces early developmental defects that result in embryonic lethality^8,9^. Environmental factors can impact both early embryogenesis, inducing phenotypes resulting in embryonic lethality as well as phenotypic changes in offspring that bypass early lethality and emerge at later developmental stages^1–4,6^. In contrast to the well-documented list of environmental maternal effects, the number of genetic maternal effects identified to induce late-stage phenotypic changes remains limited.

Approximately 70 maternal effect genes have been identified in mammals and a similar number are characterized in zebrafish^2,8– 10^. Maternal effect genes establish molecular or structural conditions that shape post-embryonic processes^9^. For example, maternal contribution of *autism susceptibility candidate 2* (*auts2*) is known to be important for offspring’s neural development. Offspring of *auts2* mutant mothers bypass embryonic lethality but display behavioral issues and differential gene expression in several brain cell types including neural crest cells and differentiating neurons^11^. *fmr1* is another maternal effect gene, where offspring of mothers mutant in *fmr1* have shorter myelin sheaths surrounding oligodendrocytes than their wildtype counterparts^12^. In *Drosophila*, maternal deposition of *germ cell-less (gcl)* is required for germ cell formation in offspring, and embryos from *gcl* mutant mothers develop somatically but lack a germline, resulting in sterility^13^. Similarly, in mice, partial loss of the *Kdm1a* gene in mothers results in abnormal behavior in the offspring once they reach adulthood like excessive scratching and digging^3^. These cases highlight how maternal gene products can have long-lasting effects in offspring, extending beyond embryogenesis and shaping critical physiological and cellular functions. However, which terminally differentiated cell populations are susceptible to genetic maternal effects remains largely unknown.

One cell type that is susceptible to late-onset environmental maternal effects is microglia, the resident immune cell of the central nervous system (CNS)^4^. Microglia have critical functions during early development, including the clearance of dead cells^14^ and neuronal circuit refinement^15^. In animal models, environmental factors during gestation like isolation and pollution agents, can cause sex-specific differences in the offspring’s microglia^4^. These offspring also display social behavior defects^4^, indicating that these phenotypic changes to microglia can impact the ability of the progeny to thrive. Immune activation in mothers has also been linked to abnormal microglia in their offspring^5^. While microglia have not yet been shown to be influenced by genetic maternal effects, we know these cells are specified early in development in the embryonic yolk sac^16,17^. Precursors in the embryonic yolk sac make both microglia and tissue-resident macrophages^18^. Recent research has shed light on how and when these two cell types are fated, but there is still much to be discovered about the genetic determinants that separate these cell types^18–20^. Whether genetic maternal effects could impact these fate decisions, and ultimately impact microglia, tissue-resident macrophages, or both, remains unanswered.

While investigating microglia in the zebrafish brain we uncovered a mutation in *sry-related HMG box gene-17* (*sox17)* that exhibits a genetic maternal effect. *sox17* is a transcription factor essential for endoderm development and maintenance of fetal hematopoietic stem cells^21,22^. As these hematopoietic stem cells differentiate, *Sox17* expression decreases, and *Sox17* mutant mice display severe hematopoietic defects and lack definitive hematopoietic stem cells^22^. Given the importance of hematopoietic stem cells in blood and immune development, *sox17* is an essential component of embryonic development, and S*ox17* null mutations are embryonic lethal^23–25^. However, sublethal *SOX17* variants in humans do not cause lethality and have been implicated in congenital anomalies of the kidneys, urinary tract, heart, and lungs^26–29^. While *Sox17* has been mostly studied in endoderm specification^23,30,31^, we uncovered that *sox17* is expressed in embryonic microglia and their yolk sac precursors. By generating a sublethal *sox17* mutation variant in zebrafish that mimics the human *SOX17* variant^28^, we identified its critical effect on microglia development that persists to later stages of development. Maternal zygotic *sox17* mutants display a reduced number of microglia compared to zygotic mutants, identifying *sox17* as a maternal effect gene critical for microglia development. Our evidence indicates that *sox17* is maternally-loaded and that restoring maternal distribution in *sox17* mutant animals can reverse the maternal effect on microglia and yolk sac cells. To determine how *sox17* could function to control microglia development, we conducted a reverse genetic screen that revealed new modifiers of developing microglia. We nominated genes for this screen that were enriched in mouse embryonic microglia from scRNA sequencing^32^. Genetic analyses with the screen hits and *sox17* mutants suggest multiple genetic pathways governing microglia abundance. Together, these results identify *sox17* as a maternal effect gene essential for microglia abundance and reveal the susceptibility of terminally-differentiated cells, like microglia, to genetic maternal effects.

## RESULTS

### *sox17* mutagenesis reduces microglia abundance in the developing brain

Previous studies have shown that severe *sox17* mutations cause embryonic lethality, so we needed to identify a sublethal variant that would allow us to assess later developmental impacts and potential maternal effects^23^. We searched the literature for *sox17* mutations that resulted in viable animals but perturbed *sox17* function. This search uncovered a mutation in humans that truncates the C-terminus of the protein and is linked to human vascular disease^28^. The location of this lesion was previously identified to impact Sox17 transcriptional targets in *Xenopus*^33^. To generate a sublethal *sox17* mutant, we injected *Tg(mrc1a:GFP)* animals at the one cell stage with Cas9 and two guide RNAs (gRNAs), one targeting each of the two exons of *sox17. Tg(mrc1a:GFP)* animals mark a subset of microglia using the *mannose receptor C, type 1a* (*mrc1a*) regulatory sequence^34^. Importantly, this microglia population has also been identified in mice during development and disease^35,36^. We stained these animals with the 4C4 antibody, which specifically labels zebrafish microglia with antigenicity to Lgals3bpb^37,38^. We scored the number of *mrc1a*^+^; 4C4^+^ cells in a 100 µm z-stack of the zebrafish midbrain (experimental imaging window unless otherwise specified) at 4 days post fertilization (dpf). T7 endonuclease assay confirmed that animals had indels near the *sox17* gRNA sites. This analysis revealed 10.44 ± 1.14 *mrc1a*^+^; 4C4^+^ cells in the brains of *sox17* crispants compared to 16.57 ± 0.95 cells in uninjected (p=0.0001) and 16.08 ± 0.90 cells in Cas9-injected controls (p=0.0007) (**Figure 1A-1B**), indicating that sublethal *sox17* perturbations can impact embryonic microglia abundance.

**Figure 1.**
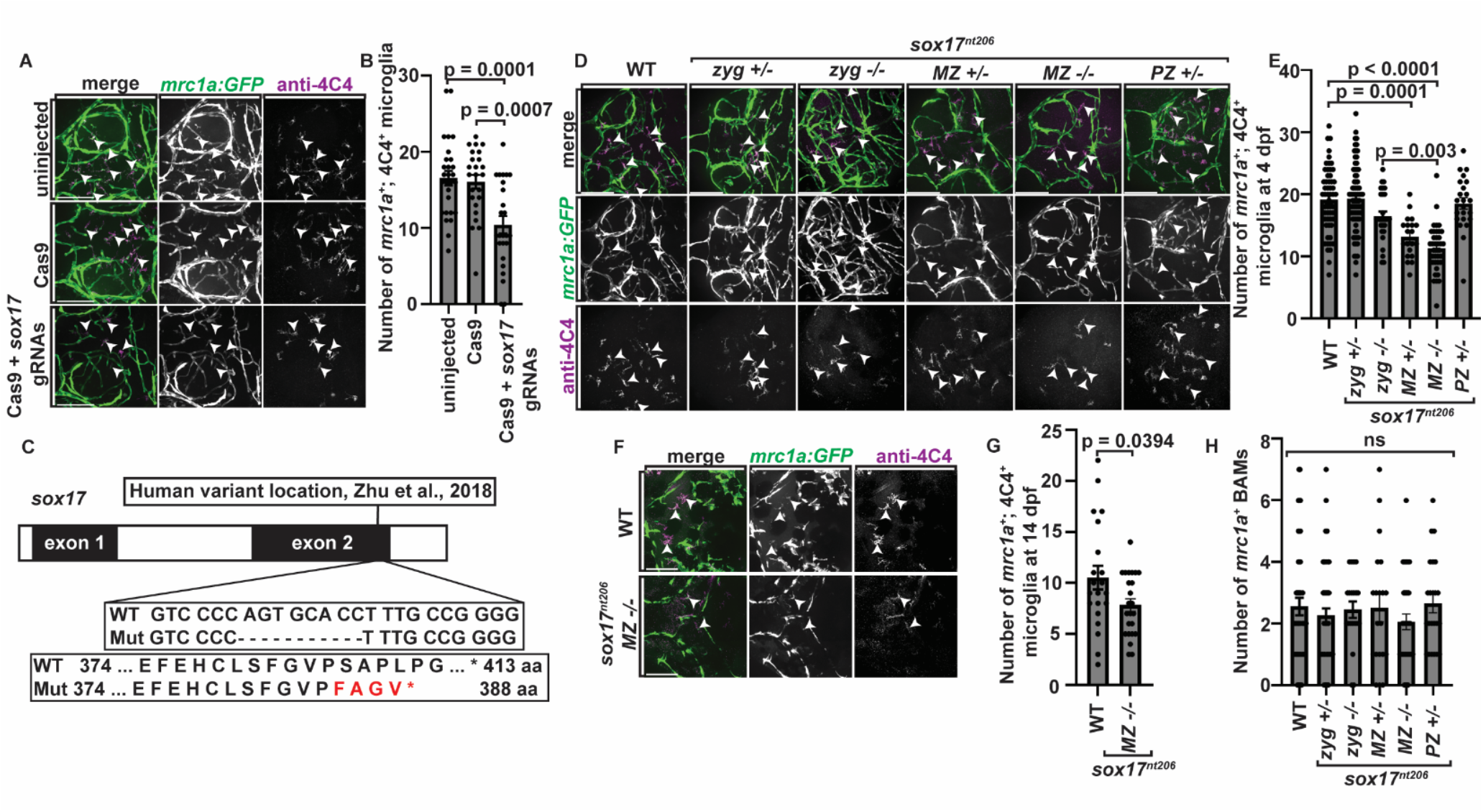
*sox17* maternal mutants have reduced microglia. **(A)** Confocal z-projections of the midbrains of 4 dpf *Tg(mrc1a:GFP)* uninjected, Cas9-injected, and Cas9 + *sox17* gRNA-injected animals stained with 4C4. **(B)** Quantification of the number of *mrc1a*^+^; 4C4^+^ microglia in the midbrains of 4 dpf *Tg(mrc1a:GFP)* uninjected (n=28), Cas9-injected (n=24), and Cas9 + *sox17* gRNA-injected (n=25) animals. **(C)** Schematic depicting the 2 exons of the *sox17* gene. Mutants have an eight-base pair deletion resulting in an early stop codon. The location of a similar human mutation variant is also shown. **(D)** Confocal z-projections of the midbrains of 4 dpf *Tg(mrc1a:GFP)* animals of various genotypes stained with 4C4. **(E)** Quantification of the number of *mrc1a*^+^; 4C4^+^ microglia in the midbrains of 4 dpf *Tg(mrc1a:GFP)* WT (n=45), *zyg*+/-(n=76), *zyg*-/- (n=28), *MZ*+/- (n=20), *MZ*-/- (n=37), and *PZ*+/- (n=23) animals. **(F)** Confocal z-projections of the midbrains of 14 dpf *Tg(mrc1a:GFP)* WT and *MZ*-/- animals stained with 4C4. **(G)** Quantification of the number of *mrc1a*^+^; 4C4^+^ microglia in the midbrains of 14 dpf *Tg(mrc1a:GFP)* WT (n=21) and *MZ*-/- (n=24) animals. **(H)** Quantification of the number of *mrc1a*^+^ BAMs in 4 dpf *Tg(mrc1a:GFP)* WT (n=45), *zyg*+/-(n=57), *zyg*-/- (n=20), *MZ*+/- (n=20), *MZ*-/- (n=34), and *PZ*+/- (n=23) animals. White arrowheads point to *mrc1a*^+^; 4C4^+^ microglia (A, D, and F). Scale bars equal 100 µm (A, D, and F). Error bars denote ± SEM (B, E, G, and H). The raw data for each group and statistical analyses are displayed in **supplemental table 2**.

### Maternal contribution of *sox17* regulates embryonic microglia abundance

To further investigate the role of *sox17* in microglia development, we generated a stable mutant using our CRISPR guide that targeted exon 2 of *sox17*. These *sox17^nt206^* animals have an eight-base pair deletion resulting in a frameshift and early stop codon in the C-terminus (**Figure 1C**). The mutant protein is predicted to be 388 amino acids compared to wildtype protein that is 413 amino acids. We generated zygotic *sox17* mutants by crossing *Tg(mrc1a:GFP); sox17^nt206+/-^* animals to each other, giving rise to wildtype, zygotic heterozygous (*zyg*+/-), and zygotic homozygous animals (*zyg*-/-). To determine the effect of *sox17* mutation on microglia development, we stained *Tg(mrc1a:GFP)* animals for 4C4 at 4 dpf and scored the number of *mrc1a*^+^; 4C4^+^ microglia in the midbrain. Quantifications demonstrated that *zyg*-/- animals had 16.46 ± 0.76 microglia compared to *zyg*+/- and wildtype siblings that had 19.32 ± 0.61 (p=0.0640) and 20.18 ± 0.73 (p=0.0129) microglia, respectively (**Figure 1D-1E**). These results are consistent with those of the *sox17* crispants and indicate that loss of *sox17* influences microglia development.

In efforts to expand generations of *sox17* mutant animals, we discovered that *zyg*-/- mutant animals survive to adulthood. Crossing the F1 generation generated 25 (52%) wildtype, 21 (43%) heterozygous, and 2 (4.1%) homozygous animals that survived to adulthood (**Figure S1A-S1B**). In a homozygous incross of F2 generation adults, several tanks of homozygous mutants survived to adulthood (data not shown), confirming that the *sox17* allele we generated is sublethal. This sublethal feature allowed us to determine the potential for maternal and paternal genetic effects. We tested this potential by comparing microglia abundance in progeny from homozygous parents. We crossed *zyg*-/- animals to generate embryos that received no contribution of wildtype *sox17* from either parent, termed maternal zygotic homozygotes (*MZ*-/-). Staining with 4C4 and quantifying the abundance of microglia at 4 dpf in the midbrains of *MZ*-/- animals showed 11.30 ± 0.62 *mrc1a*^+^; 4C4^+^ cells, revealing a significant decrease in microglia number compared to *zyg*-/- animals (16.46 ± 0.76 microglia, p=0.0002), whose parents were heterozygotes for mutated *sox17* (**Figure 1D-1E**). These data indicate that homozygous offspring, generated from homozygous mutant parents, have significantly fewer microglia than homozygous offspring generated from heterozygous mutant parents, suggesting a maternal or paternal genetic effect impacting microglia abundance.

To understand if there was a genetic maternal effect of *sox17* on microglia abundance, we crossed maternal *sox17*^-/-^ animals to wildtype paternal animals. This mating generated maternal zygotic heterozygous (*MZ*+/-) progeny due to the lack of maternally-contributed *sox17*. Staining *MZ*+/- animals with 4C4 and quantifying the abundance of microglia in the midbrain at 4 dpf showed 13.10 ± 0.73 *mrc1a*^+^; 4C4^+^ cells, significantly fewer than wildtype animals (20.18 ± 0.73 microglia, p<0.0001) and phenocopying *MZ*-/- microglia numbers (11.30 ± 0.62 microglia, p=0.7262) (**Figure 1D-1E**). These data show that progeny generated from *sox17*^-/-^ mothers have a drastic reduction in microglia, regardless of if the progeny genotype is *MZ*-/- or *MZ*+/-. This result reveals that a lack of maternal contribution of wildtype *sox17* is sufficient to reduce microglia number.

To test if the decrease in microglia abundance was specific to a lack of maternally-contributed *sox17*, we crossed maternal wildtype animals to paternal *sox17*^-/-^ animals to create paternal zygotic heterozygous (*PZ*+/-) mutants. Scoring *mrc1a*^+^; 4C4^+^ microglia abundance at 4 dpf showed 18.52 ± 0.92 microglia in the midbrains of *PZ*+/- animals, a number consistent with wildtype (20.18 ± 0.73 microglia, p=0.7307) and *zyg*+/- animals (19.32 ± 0.61 microglia, p=0.9794) (**Figure 1D-1E**). These data support the idea that maternally-contributed *sox17* is required for proper embryonic microglia abundance. While we considered that *sox17^nt206^* animals could have an off-target mutation that was generated during CRISPR mutagenesis, the clear phenotype of *MZ*+/- mutants generated from paternal wildtype animals is inconsistent with that explanation. The normal abundance of microglia in *PZ*+/- and *zyg*+/- also ruled out the potential of the *sox17*^*nt206*-/-^ allele functioning as a dominant negative. The simplest explanation of this data is that the maternal contribution of *sox17* is critical for microglia development, and thus indicates microglia are susceptible to a genetic maternal effect. We focused on this maternal effect moving forward by investigating the maternal zygotic homozygous (*MZ*-/-) progeny.

We next scored the effect of maternal *sox17* on total 4C4^+^ microglia in the brain, regardless of *mrc1a*^+^ expression, to capture all subsets of microglia. Compared to wildtype animals which had 25.07 ± 0.72 4C4^+^ microglia, we scored 26.17 ± 0.74, 24.42 ± 1.03, 24.1 ± 0.83, and 26.52 ± 1.27 4C4^+^ microglia in the midbrains of *zyg*+/- (p=0.8677), *zyg*-/- (p=0.9959), *MZ*+/- (p=0.9841), and *PZ*+/- (p=0.8949) animals, respectively (**Figure S1C**). We scored 15.49 ± 0.60 4C4^+^ microglia in *MZ*-/- animals, a significant decrease compared to wildtype animals (p<0.0001) (**Figure S1C**). The normal 4C4^+^ microglia abundance in *MZ*+/- animals indicates that the maternal requirement of *sox17* on microglia is likely unique to the *mrc1a*^+^subpopulation of microglia, although microglia number is reduced in *MZ*-/- animals, regardless of *mrc1a* expression.

### Microglia abundance in *sox17 MZ*-/- mutants is reduced until at least 14 dpf

Genetic maternal effects typically induce phenotypes in early embryogenesis, but the maternal effect of *sox17* on microglia is apparent at 4 dpf, a time point beyond embryogenesis in zebrafish^9,39^. Because of this, we asked if the genetic maternal effect of *sox17* continues to impact microglia abundance at later developmental stages. To assess the maternal effect of *sox17* on microglia throughout development, we grew wildtype and *MZ-/-Tg(mrc1a:GFP)* animals to 14 dpf. Staining with 4C4 and scoring microglia number revealed 10.52 ± 1.16 *mrc1a*^+^; 4C4^+^ microglia in wildtype animals, compared to 7.83 ± 0.62 in *MZ*-/- mutants (p=0.0394) (**Figure 1F-1G**), indicating a persistence in microglia reduction in *MZ*-/- mutants to at least 14 dpf. Similarly, we observed 15.24 ± 10.42 4C4^+^ microglia in the brains of wildtype animals compared to 10.42 ± 0.91 microglia in *MZ*-/- mutants (p=0.0048) (**Figure S1D**), indicating that the effect of *sox17* on microglia abundance, regardless of *mrc1a*^+^ expression, persists until at least 14 dpf. These data further reveal this unique, late-stage genetic maternal effect of *sox17* on microglia abundance.

### *sox17* mutation does not impact BAMs or brain vasculature

Given that maternal *sox17* is important for *mrc1a*^+^ microglia abundance in the brain at 4 dpf, and microglia and macrophages share a common *Mrc1*^+^ precursor, we next investigated the impact of *sox17* on the brain-associated macrophage population^18^. Border associated macrophages (BAMs) reside at CNS borders like the meninges and perivascular spaces, and rely on many of the same genetic determinants as microglia^18,40,41^. Microglia and BAMs were originally shown to be fated separately in the embryonic yolk sac, but recent studies have revealed they share a common *Mrc1*^+^ progenitor^18,19^. Given this, we asked if *sox17* mutation also impacted the BAM population in 4 dpf animals. Wildtype and *MZ-/-Tg(mrc1a:GFP)* animals were grown to 4 dpf and stained with 4C4 to distinguish 4C4^+^ microglia from 4C4^-^ BAMs. Analysis revealed no difference in the number of BAMs between wildtype (4.07 ± 0.30 BAMs) and *zyg*+/- (3.95 ± 0.23 BAMs, p>0.9999), *zyg*-/- (3.45 ± 0.43 BAMs, p>0.9999), *MZ*+/- (2. 95 ± 0.46 BAMs, p=0.5246), *MZ*-/- (3.24 ± 0.30 BAMs, p=0.6142), or *PZ*+/- (3.09 ± 0.35 BAMs, p=0.6598) animals (**Figure 1H**). These data indicate that *sox17* mutation does not cause widespread abnormalities in myeloid cells but rather specifically impacts microglia.

We next considered the hypothesis that the maternal effect of *sox17* on microglia could be driven by patterning defects of other tissues. *sox17* is a regulator of vascular development, and microglia are known to be impacted by defects in vasculature^30,34,42,43^. As *mrc1a* also marks lymphatic and venous vasculature^44^, we used *Tg(mrc1a:GFP)* animals to test if there was a maternal effect of *sox17* on vascular patterning. We scored patterning of the vasculature in and around the midbrain by several different measures. We first analyzed the length of the midcerebral vein (**Figure S1E, label F**) and the mesencephalic vessels (**Figure S1E, label G**). We also scored the sum of the area of the two vascular loops dorsal to the midbrain (**Figure S1E, label H**), as well as the percent of the image covered by GFP^+^ signal to assess overall vascular abundance (**Figure S1E**)^45,46^. Because some 4 dpf zebrafish exhibit autofluorescence in the skin, we thresholded the images when quantifying the percent of the image covered by signal to exclude any images that had abundant signal coverage due to autofluorescence (**Figure S1E**). Compared to wildtype animals, in *zyg+/-, zyg*-/-, or *MZ*-/- animals there were no differences detected in the length of the midcerebral vein (**Figure S1F)**, the length of the mesencephalic vessel (**Figure S1G**), the sum of the area of the vascular loops (**Figure S1H**), or the percent of the image covered by GFP+ signal (**Figure S1F-I**). These results suggest that *sox17* mutation does not impair vascular patterning and therefore the reduction in microglia abundance in *MZ*-/- animals is not likely to be due to vascular patterning defects.

### *mrc1a*^+^ microglia express *sox17*

To next gain insight into how *sox17* could impact microglia, we examined the spatial expression pattern of *sox17* under normal conditions. We used *Tg(sox17:H2A-mCherry)* animals that use regulatory regions of the *sox17* promoter to drive expression of nuclear-localized mCherry. Scoring the percent of *mrc1a*^+^; 4C4^+^ cells with mCherry^+^ expression in 4 dpf wildtype *Tg(mrc1a:GFP); Tg(sox17:H2A-mCherry)* animals revealed 4.5 ± 1.45% of *mrc1a*^+^; 4C4^+^ microglia expressing *sox17* in wildtype animals (**Figure 2A-2B**). These data identify that *sox17* is expressed in *mrc1a*^+^ microglia, supporting the idea that *sox17* is required for proper microglia development. We also scored *sox17* expression in *pu1*^+^ microglia using wildtype *Tg(pu1:Gal4; UAS:GFP); Tg(sox17:H2A-mCherry)* fish and scored 7.8 ± 3.16% of *pu1*^+^; 4C4^+^ microglia expressing *sox17* (**Figure S2A-S2B**). Coupled with our data above that *sox17 MZ*-/- mutants have a reduction in total 4C4^+^ microglia, these data suggest that *sox17* impacts multiple microglia subpopulations in the developing brain.

**Figure 2.**
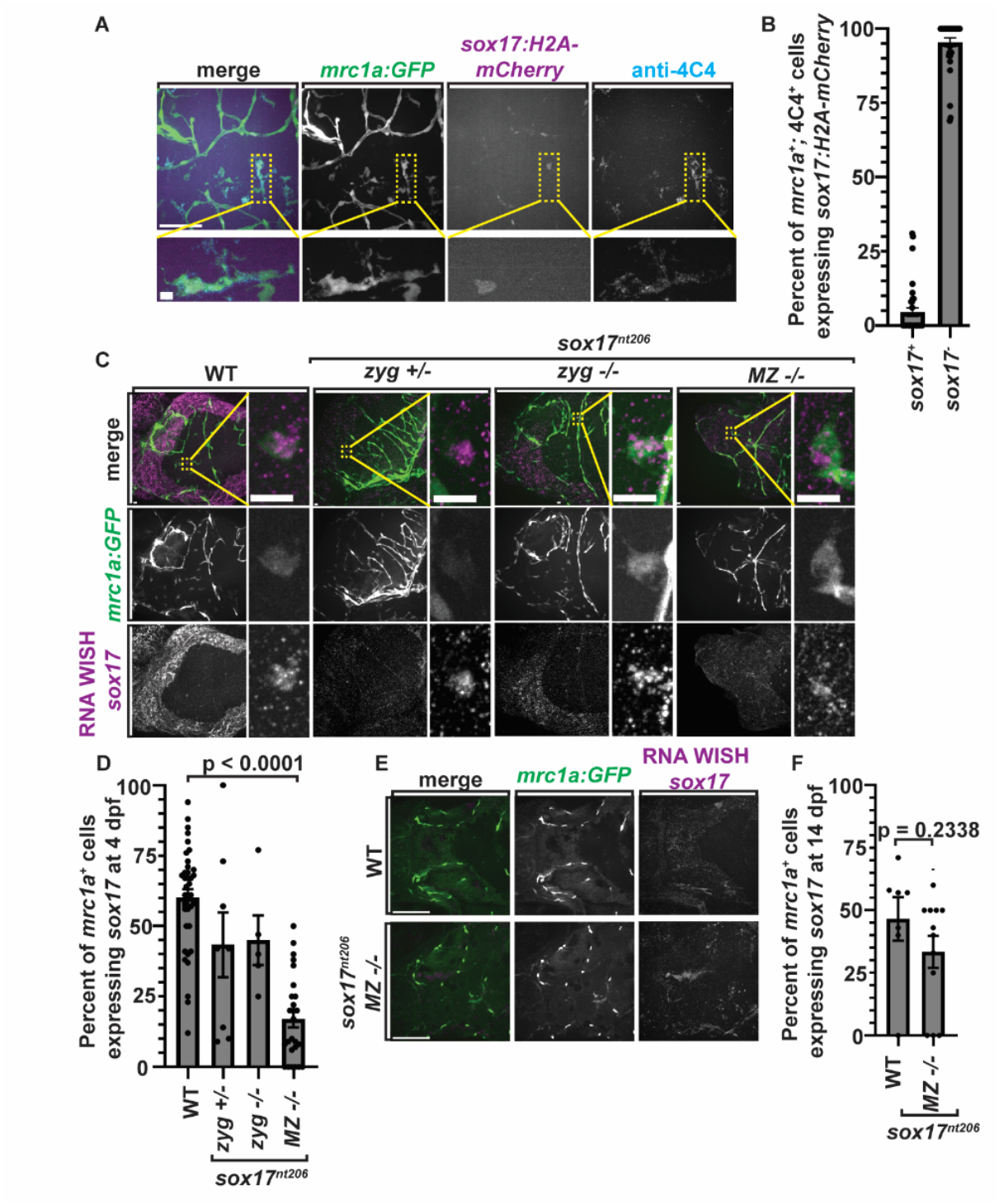
*sox17* is expressed in microglia. **(A)** Confocal z-projections of the midbrains of 4 dpf *Tg(mrc1a:GFP); Tg(sox17:H2A-mCherry)* animals stained with 4C4. Yellow boxes/insets show a single cell co-expressing *mrc1a:GFP* and *sox17:H2A-mCherry.* **(B)** Quantification of the percent of *mrc1a*^+^; 4C4^+^ microglia that are *sox17:H2A-mCherry*^+^ or *sox17:H2A-mCherry*^-^ in the midbrain of 4 dpf *Tg(mrc1a:GFP); Tg(sox17:H2A-mCherry)* animals (n=35). **(C)** Confocal z-projections of the midbrains of 4 dpf *Tg(mrc1a:GFP)* animals of various genotypes stained with WISH HCR against *sox17.* Yellow boxes/insets show a single cell co-expressing *mrc1a:GFP* and *sox17.* **(D)** Quantification of the percent of *mrc1a*^+^ cells labeled with WISH HCR against *sox17* in the midbrain of 4 dpf *Tg(mrc1a:GFP)* WT (n=37), *zyg*+/-(n=8), *zyg*-/- (n=5), and *MZ*-/- (n=26) animals. **(E)** Confocal z-projections of the midbrains of 14 dpf *Tg(mrc1a:GFP)* animals of various genotypes stained with WISH HCR against *sox17*. **(F)** Quantification of the percent of *mrc1a*^+^ cells that labeled with HCR against *sox17* in the midbrain of 14 dpf *Tg(mrc1a:GFP)* WT (n=7), and *MZ*-/- (n=12) animals. Yellow inset scale bars equal 10 µm (A and C insets). All other scale bars equal 100 µm (A, C, and E). Error bars denote ± SEM (B, D, and F). The raw data for each group and statistical analyses are displayed in **supplemental table 2**.

### *sox17* mutants have a reduction in *sox17* transcript

To understand the mechanism of this maternal effect of *sox17* and provide a complementary approach to determine *sox17* expression, we next asked if *sox17* mutants display reduced *sox17* transcript levels, and how transcript levels differed between genotypes using *in situ* hybridized chain reaction (HCR) on whole mount embryos. We first assayed *sox17* transcript expression at 4 dpf when the number of microglia is reduced. We assayed *sox17* transcript in wildtype, *zyg+/-, zyg*-/-, and *MZ*-/- genotypes. Because we know that the various genotypes have differing numbers of *mrc1a*^+^ cells, the percent of total *mrc1a*^+^ cells expressing *sox17* transcript was scored. Imaging of the midbrain revealed 60.14 ± 3.03% *mrc1a*^+^ cells expressing *sox17* transcript in wildtype animals, compared to 43.38 ± 11.52% in *zyg*+/- animals (p=0.3349), 45.00 ± 8.76% in *zyg*-/- animals (p=0.6950), and 17.04 ± 2.98% in *MZ*-/- mutants (p<0.0001) (**Figure 2C-2D**). *MZ*-/- animals have a significant reduction in the total number of *mrc1a*^+^ cells that express *sox17* transcript at 4 dpf. These data reveal a fascinating finding that a lack of maternally-contributed *sox17* in *MZ*-/- mutants reduces *sox17* transcript expression in the offspring - an impact that persists far past the typical timing of genetic maternal effects.

To further probe this effect on transcript expression, we next asked if the maternal effect on *sox17* transcription was still detectable at 14 dpf, a time point when the reduction of microglia is still present (**Figure 1G)**. Scoring the percent of *mrc1a*^+^ cells expressing *sox17* transcript in 14 dpf wildtype and *MZ-/-Tg(mrc1a:GFP)* animals revealed no difference between wildtype (46.57 ± 8.69%) and *MZ*-/- (33.42 ± 6.37%) animals (p=0.2338) (**Figure 2E-2F**). These data suggest that the maternal effect of *sox17* on transcript abundance is resolved by 14 dpf, even though there is still a reduction in the number of *mrc1a*^+^ microglia at this time point. These results are consistent with the idea that *sox17* exhibits a significant genetic maternal effect on *sox17* transcript levels that persists through 4 dpf.

### *sox17* is maternally contributed to offspring

Genetic maternal effects depend on the contribution of maternally distributed RNA or protein^47^. Maternal RNA is contributed to embryos throughout early embryogenesis, prior to zygotic genome activation^48^. Zygotic genome activation begins roughly 2-4 hours post fertilization (hpf) in zebrafish^49^. After 4 hpf, maternal transcripts decrease as zygotic transcripts increase^50^. Since the most severe phenotypes in reduction of microglia and *sox17* transcript are seen in *MZ*-/- mutants, we next sought to confirm that *sox17* is maternally contributed to progeny during early embryogenesis. To do this we collected RNA from wildtype and *MZ-/-Tg(mrc1a:GFP)* animals at 1 hpf, when zygotic genome activation has not yet begun and thus any transcript would be maternally-contributed. Isolation of RNA and amplification of its cDNA for a ∼300 bp region of the *sox17* mRNA showed a clear product at 1 hpf, indicating that *sox17* is maternally-loaded (**Figure 3A**, full gel image **Figure S3C**). Given that our *sox17* mutation results in a minimally truncated protein, we would expect protein expression or function, and not RNA levels, to be impacted by mutation. These data confirm that *sox17* is maternally-loaded in the developing embryo.

**Figure 3.**
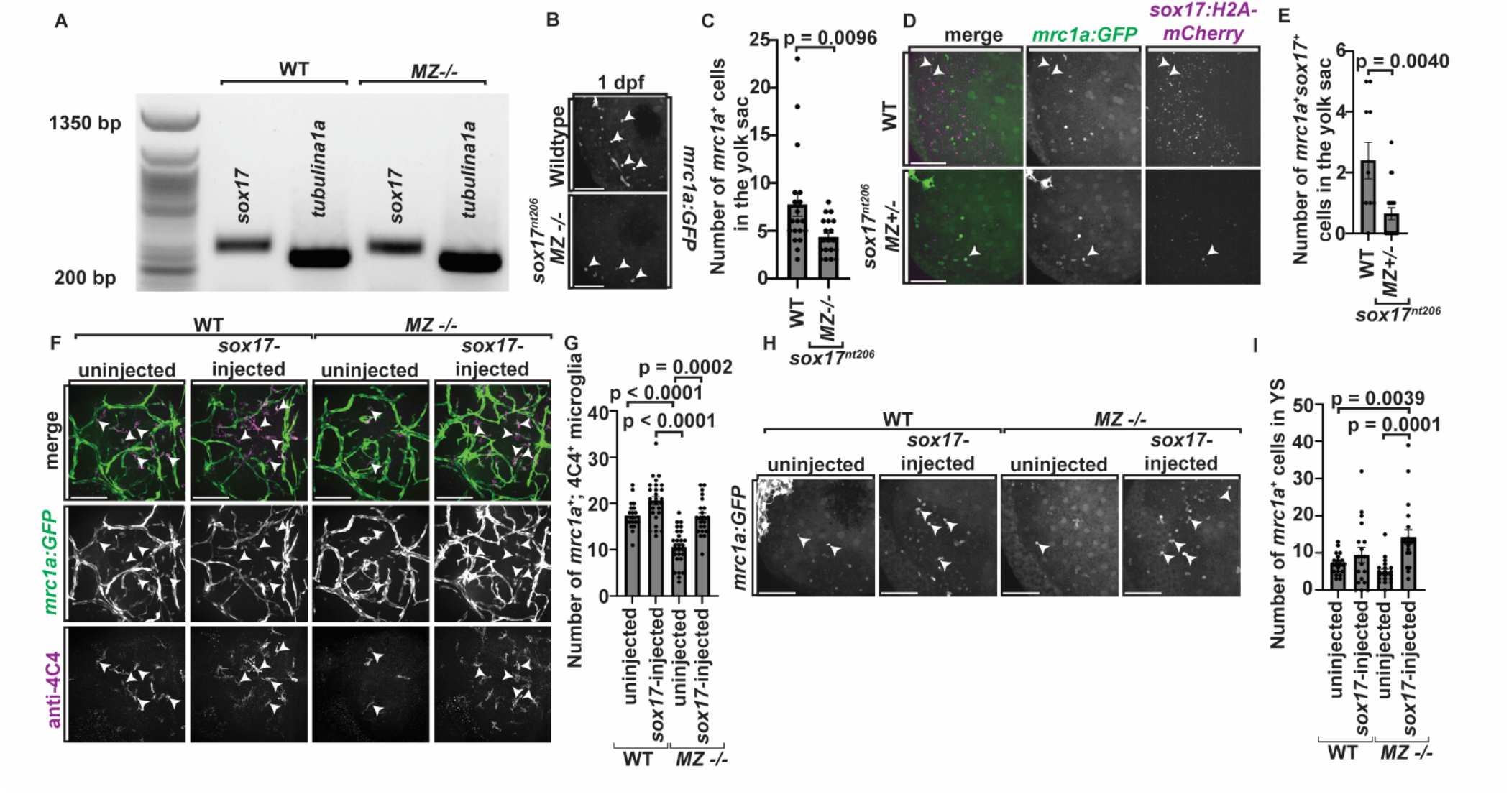
*sox17* is maternally loaded and important for yolk sac microglia precursors. **(A)** Agarose gel showing PCR on cDNA for *sox17* and *tubulina1a* in 1 hpf WT and *MZ*-/- animals. **(B)** Confocal z-projections of the yolk sacs of 1 dpf *Tg(mrc1a:GFP)* WT and *MZ*-/- animals. White arrowheads point to *mrc1a*^+^ cells. **(C)** Quantification of the number of *mrc1a*^+^ cells in the yolk sac of 1 dpf *Tg(mrc1a:GFP)* WT (n=19) and *MZ*-/- (n=19) animals. **(D)** Confocal z-projections of the yolk sacs of 1 dpf *Tg(mrc1a:GFP); Tg(sox17:H2A-mCherry)* WT and *MZ*+/- animals. White arrowheads point to *mrc1a^+^; sox17*^+^ cells. **(E)** Quantification of the number of *mrc1a^+^; sox17*^+^ cells in the yolk sac of 1 dpf *Tg(mrc1a:GFP); Tg(sox17:H2A-mCherry)* WT (n=10) and *MZ*+/- (n=20) animals. **(F)** Confocal z-projections of the midbrains of 4 dpf *Tg(mrc1a:GFP)* uninjected or *sox17-P2A-mCherry-pA* RNA-injected WT and *MZ*-/- animals stained with 4C4. White arrowheads point to *mrc1a*^+^; 4C4^+^ microglia. **(G)** Quantification of the number of *mrc1a*^+^; 4C4^+^ microglia in the midbrain of 4 dpf *Tg(mrc1a:GFP)* uninjected WT (n=19), injected WT (n=27), uninjected *MZ*-/- (n=28), and injected *MZ*-/- (n=22) animals. **(H)** Confocal z-projections of the yolk sacs of 1 dpf *Tg(mrc1a:GFP)* uninjected or *sox17-P2A-mCherry-pA* RNA-injected WT and *MZ-/-Tg(mrc1a:GFP)* animals. White arrowheads point to *mrc1a*^+^ cells. **(I)** Quantification of the number of *mrc1a*^+^ cells in the yolk sac of 1 dpf *Tg(mrc1a:GFP)* uninjected WT (n=20), injected WT (n=18), uninjected *MZ*-/- (n=20), and injected *MZ*-/- (n=20) animals. Scale bars equal 100 µm (B, D, F, and H). Error bars denote ± SEM (C, E, G, and I). The raw data for each group and statistical analyses are displayed in **supplemental table 2**.

### *sox17* mutants have fewer microglia precursors in the yolk sac

Knowing that *sox17* is maternally-loaded, we next investigated the developmental stage at which the maternal effect on *mrc1a*^+^ cells manifests. The yolk sac is present at fertilization in zebrafish and is therefore well-positioned to be impacted by maternal effect genes^51^. Recent work supports the idea that *mrc1a*^+^ cells present in the yolk sac at 1 dpf colonize the developing brain^34^. Because microglia originate in the embryonic yolk sac, we investigated microglia precursors in the yolk sac to begin to dissect the mechanism of the genetic maternal effect of *sox17* on microglia abundance^16,17^. To ask if *sox17* could have a maternal effect on microglia progenitors in the yolk sac we scored the number of *mrc1a*^+^ cells in the yolk sac of 1 dpf wildtype and *MZ-/-Tg(mrc1a:GFP)* animals, which revealed 7.74 ± 1.21 cells in the yolk sacs of wildtype animals compared to 4.32 ± 0.44 cells in the yolk sacs of *MZ*-/- animals (p=0.0096) (**Figure 3B-3C**), and consistent with the hypothesis that *sox17* plays an important role in *mrc1a*^+^ cell production in the yolk sac.

### *mrc1a*^+^ cells in the yolk sac express *sox17*

To further understand the requirement of *sox17* in production of *mrc1a*^+^ yolk sac cells, we asked if *mrc1a*^+^ yolk sac cells express *sox17*. Wildtype and *MZ-/-Tg(mrc1a:GFP)* animals were crossed to *Tg(sox17:H2A-mCherry)* animals to generate wildtype and *MZ+/-Tg(mrc1a:GFP); Tg(sox17:H2A-mCherry)* animals. We scored the number of double positive *mrc1a*^+^; *sox17*^+^ cells in the yolk sacs at 1 dpf and uncovered 2.4 ± 0.60 *mrc1a*^+^; *sox17*^+^ cells in the yolk sacs of wildtype animals, compared to 0.65 ± 0.20 *mrc1a*^+^; *sox17*^+^ cells in the yolk sacs of *MZ*+/- mutants (p=0.0040) (**Figure 3D-3E**). These data reveal that *mrc1a*^+^ cells in the yolk sac express *sox17*, supporting the hypothesis that

*mrc1a*^+^ yolk sac cells are likely influenced by maternally-contributed *sox17*. These data also confirmed that, just as we measured with *mrc1a*^+^; 4C4^+^ microglia, even *MZ*+/- mutants display a reduction in *mrc1a*^+^ yolk sac cells, suggesting that even one paternally-contributed wildtype copy of *sox17* is not sufficient to drive proper development of *mrc1a*^+^ cells in the yolk sac.

### *sox17* rescue restores abundance of *mrc1a*^+^ microglia in the brain and *mrc1a*^+^ cells in the yolk sac

Maternal RNA functions in the embryo until zygotic genome activation^48,50^. We hypothesized that if *sox17* functioned as a maternally distributed product, that restoring *sox17* mRNA before zygotic genome activation should restore microglia numbers in *MZ*-/- progeny. To do this, cDNA corresponding to the coding sequence of *sox17* was fused with a P2A-mCherry-pA tag to create a *sox17-P2A-mCherry-pA* construct. This construct was *in vitro* transcribed, and the RNA was injected into the one cell stage of wildtype and *MZ-/-Tg(mrc1a:GFP)* animals. This time precedes zygotic genome activation^49^ and introduces *sox17-P2A-mCherry-pA* RNA at a time when only maternal transcripts are present. At 1 dpf, animals showed no gross morphological defects and were screened for mCherry^+^ expression (**Figure S3A**) and then grown to 4 dpf. We assayed the effect of *sox17* rescue on microglia abundance in the brain at 4 dpf in both wildtype and *MZ*-/- animals that were uninjected or injected with *sox17-P2A-mCherry-pA* RNA. In wildtype animals where maternal contributions of *sox17* were not disrupted, injection of *sox17-P2A-mCherry-pA* RNA had no effect on the number of *mrc1a*^+^; 4C4^+^ microglia (20.63 ± 0.88 microglia) in the brain at 4 dpf compared to uninjected wildtype animals (17.42 ± 0.70 microglia, p=0.4388) (**Figure 3F-3G**). We also observed no impact on the number of total 4C4^+^ microglia in wildtype RNA-injected animals (24.70 ± 1.04 cells) compared to uninjected controls (24.32 ± 0.78, p>0.9999) (**Figure S3B**). In uninjected *MZ*-/- animals we observed 10.61 ± 0.73 *mrc1a*^+^; 4C4^+^ microglia and 15.57 ± 1.16 total 4C4^+^ cells (**Figure 3F-3G, Figure 3SB**). Injecting *sox17-P2A-mCherry-pA* RNA into *MZ*-/- animals resulted in 17.14 ± 0.83 *mrc1a*^+^; 4C4^+^ microglia and 23.23 ± 0.96 total 4C4^+^ microglia density in the brain, consistent with wildtype numbers (*mrc1a*^+^; 4C4^+^ microglia: p>0.9999, total 4C4^+^ microglia: p>0.9999) (**Figure 3F-3G, Figure 3SB**). These data support the hypothesis that mimicking maternal contribution of *sox17* RNA is sufficient to restore *mrc1a*^+^; 4C4^+^, as well as total 4C4^+^, microglia abundance in the brains of *sox17 MZ*-/- mutants, and suggests that *sox17* maternal contribution is critical for embryonic microglia development.

Because we observed fewer *mrc1a*^+^ cells in the yolk sacs of *sox17 MZ*-/- mutants compared to wildtype animals (**Figure 3C**), we also asked if *sox17* rescue could restore *mrc1a*^+^ cell numbers in the yolk sac. Similarly to above, wildtype and *MZ*-/- animals were injected with the *sox17-P2A-mCherry-pA* RNA construct at the one cell stage and screened for mCherry^+^ signal at 1 dpf. Uninjected animals were used as controls. The yolk sacs of animals were imaged at 1 dpf and the number of *mrc1a*^+^ cells was scored. Injection of *sox17-P2A-mCherry-pA* RNA in wildtype animals resulted in 9.44 ± 2.08 *mrc1a*^+^ cells in the yolk sac, compared to 7.05 ± 0.63 in the yolk sac cells of wildtype, uninjected animals (p=0.6695) (**Figure 3H-3I**). In contrast, injection of *sox17-P2A-mCherry-pA* in *MZ*-/- mutants resulted in 14.3 ± 1.96 *mrc1a*+ cells in the yolk sac, a significant increase compared to uninjected wildtype numbers (p=0.0039) and uninjected *MZ*-/- numbers (4.95 ± 0.72 cells, p=0.0001) (**Figure 3H-3I**). These data suggest that maternally-deposited *sox17* is necessary for *mrc1a*^+^ cell development in the embryonic yolk sac and are consistent with the hypothesis that *sox17* must be present prior to zygotic genome activation for proper abundance of yolk sac cells at 1 dpf and *mrc1a*^+^ microglia in the brain at 4 dpf.

### A high-throughput CRISPR screen identifies additional genetic modifiers of microglia development

To expand our understanding of how microglia development is regulated via *sox17*, particularly in the context of the identified maternal effect, we conducted a high-throughput CRISPR screen utilizing a custom-built robot that captures confocal images of the zebrafish brain in high-throughput. To nominate genes, we analyzed the E14.5 timepoint in the Hammond et al. scRNA sequencing data, a timepoint when microglia are differentiating in the mouse brain^32,52^. This analysis identified two cell clusters of interest - an embryonic microglia cluster enriched in known microglia markers such as *P2ry12, Tmem119,* and *Mrc1,* and a macrophage/microglia cluster which was enriched in *Mrc1* and *P2ry12* (**Figure 4A-4B**)^32^. In choosing genes to screen, we focused primarily on 24 genes encoding transcription factors and cell-surface receptors. We designed gRNA pools corresponding to each gene (**Table S1**). For each gene, we designed 4 corresponding gRNAs. For genes that are duplicated in zebrafish we designed 3 gRNAs (**Table S1**). Wildtype *Tg(mrc1a:GFP)* animals were injected with gRNA pools for each gene, grown to 4 dpf, imaged on our custom-built automated confocal, and the number of *mrc1a*^+^ cells in the midbrain was scored. Uninjected and Cas9-injected animals were used as controls. Importantly, our screening microscope captures whole body, brightfield images of each animal, revealing that all animals included in the quantification had no gross morphological defects (**Figure 4C**). A volcano plot was generated where we plotted the difference of number of *mrc1a*^+^ cells in the brains of Cas9-injected and Cas9 + gRNA-injected animals at 4 dpf and the corresponding p values (**Figure 4D**). Of the 24 genes screened, 6 genes, *cmtm7, cndp2, f11r.1, gas6, irf8,* and *mpp1,* showed an initial phenotype of reduced *mrc1a*^+^ cell abundance in the brain (**Figure 4C-4E**). To next determine if these genes could have evolutionarily conserved roles in microglia development, we generated UMAPs and violin plots of their expression in mouse embryonic microglia^32^. These plots show the expression of these 6 genes in our scRNA sequencing analysis of mouse embryonic microglia and macrophage clusters. *Cmtm7* (**Figure S4A, S4F**), *Cndp2* (**Figure S4B, S4G**), *F11r.1* (**Figure S4C, S4H**), *Gas6* (**Figure S4D, S4I**), *Irf8* (**Figure S4E, S4J**), and *Mpp1* (**Figure 4F-4G**) were all enriched in mouse embryonic microglia, while only *Cmtm7, Cndp2, Gas6, Irf8,* and *Mpp1* were enriched in macrophages^32^. In summary, our genetic screen identified 6 novel candidate regulators of *mrc1a*^+^ cell development in the developing brain and opens the possibility for unidentified genetic pathways governing microglia abundance.

**Figure 4.**
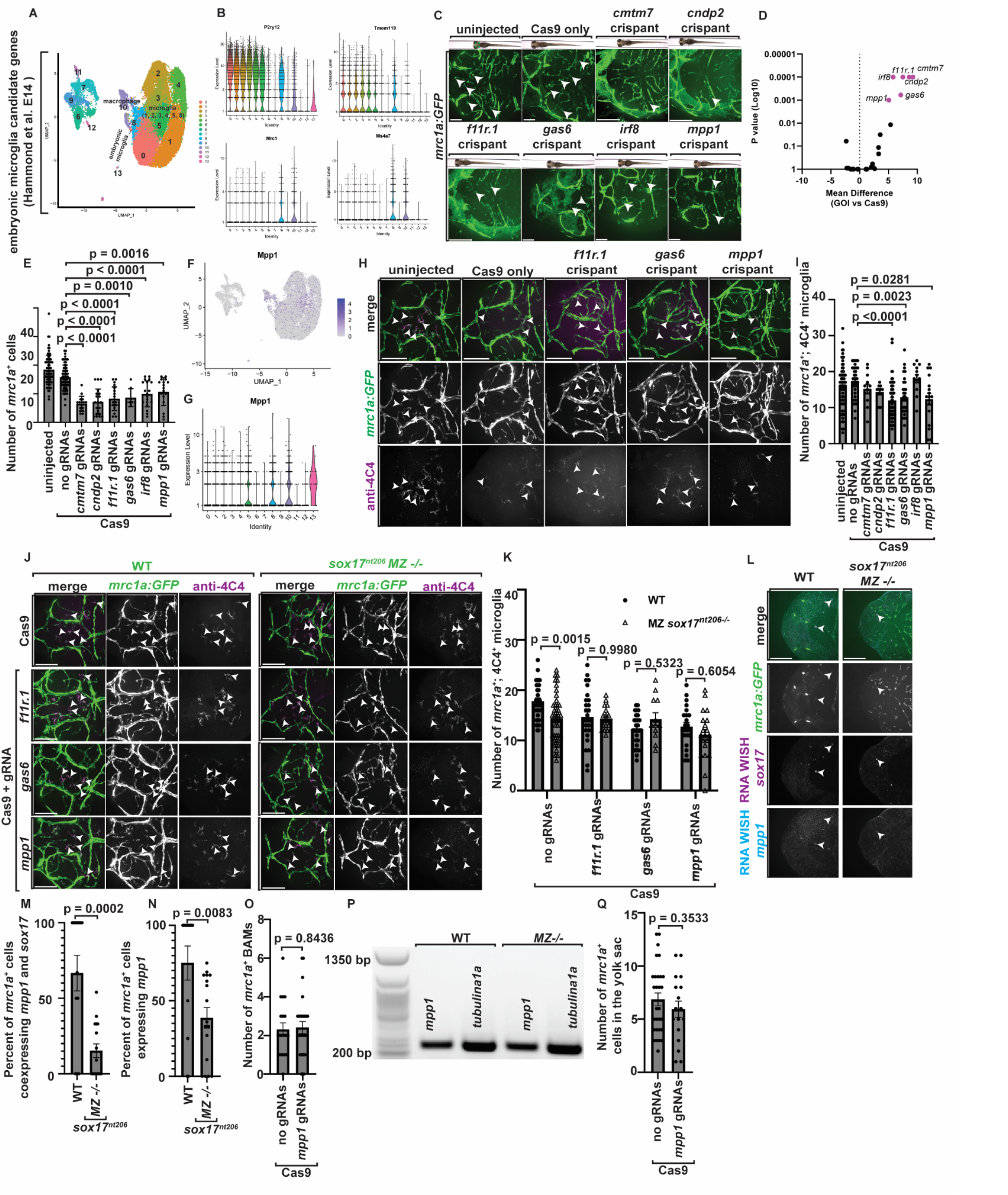
Genetic interrogation reveals genetic modifiers of microglia that coordinate with *sox17*. **(A)** UMAP representing all 13 clusters identified in our reanalysis of the Hammond et al., 2014 dataset^32^. **(B)** Violin plots of *P2ry12, Tmem119, Mrc1, and Ms4a7* across all 13 clusters shown in the UMAP^32^. **(C)** Confocal z-projections of the midbrains of uninjected, Cas9-injected, or Cas9 + gRNA pool-injected 4 dpf *Tg(mrc1a:GFP)* animals. White arrowheads point to *mrc1a*^+^ cells. **(D)** Volcano plot representing the mean difference in the number of *mrc1a*^+^ cells in the brains of Cas9-injected and Cas9+ gRNA-injected 4 dpf *Tg(mrc1a:GFP)* animals. Genes represented with pink dots displayed a significant reduction in cell number (p≤0.0010). **(E)** Quantification of the number of *mrc1a*^+^ cells in 4 dpf *Tg(mrc1a:GFP)* uninjected (n=72), Cas9-injected (n=63), Cas9 + *cmtm7* gRNA pool-(n=16), Cas9 + *cndp2* gRNA pool-(n=20), Cas9 + *f11r.1* gRNA pool-(n=19), Cas9 + *gas6* gRNA pool-(n=8), Cas9 + *irf8* gRNA pool-(n=19), and Cas9 + *mpp1* gRNA pool-injected (n=17) animals. White arrowheads point to *mrc1a*^+^ cells**. (F)** UMAP and **(G)** violin plot of *Mpp1* expression in E14 mouse microglia^32^. **(H)** Confocal z-projections of the midbrains of 4 dpf *Tg(mrc1a:GFP)* uninjected, Cas9-injected, Cas9 + *f11r.1* gRNA pool-, Cas9 + *gas6* gRNA pool-, and Cas9 + *mpp1* gRNA pool-injected animals stained with 4C4. White arrowheads point to *mrc1a*^+^; 4C4^+^ microglia. **(I)** Quantification of the number of *mrc1a*^+^; 4C4^+^ microglia in the midbrain of 4 dpf *Tg(mrc1a:GFP)* uninjected (n=66), Cas9-injected (n=38), Cas9 + *cmtm7* gRNA pool-(n=56), Cas9 + *cndp2* gRNA pool-(n=17), Cas9 + *f11r.1* gRNA pool-(n=32), Cas9 + *gas6* gRNA pool-(n=12), Cas9 + *irf8* gRNA pool-(n=10), and Cas9 + *mpp1* gRNA pool-injected (n=13) animals. **(J)** Confocal z-projections of the midbrains of 4 dpf *Tg(mrc1a:GFP)* WT and *MZ*-/- animals injected with Cas9, Cas9 + *f11r.1* gRNA pool, Cas9 + *gas6* gRNA pool, or Cas9 + *mpp1* gRNA pool stained with 4C4. White arrowheads point to *mrc1a*^+^; 4C4^+^ microglia. **(K)** Quantification of the number of *mrc1a*^+^; 4C4^+^ microglia in the midbrain of 4 dpf *Tg(mrc1a:GFP)* wildtype animals injected with Cas9 (n=39), Cas9 + *f11r.1* gRNA pool (n=28), Cas9 + *gas6* gRNA pool (n=28), or Cas9 + *mpp1* gRNA pool (n=23) and 4 dpf *Tg(mrc1a:GFP) MZ*-/- animals injected with Cas9 (n=42), Cas9 + *f11r.1* gRNA pool (n=18), Cas9 + *gas6* gRNA pool (n=12), and 4 dpf Cas9 + *mpp1* gRNA pool (n=20). **(L)** Confocal z-projections of the midbrains of 4 dpf *Tg(mrc1a:GFP)* animals stained with WISH HCR probes against *sox17* and *mpp1*. White arrowheads point to *mrc1a^+^; sox17^+^; mpp1*^+^ cells. **(M)** Quantification of the percent of *mrc1a*^+^ cells co-expressing *sox17* and *mpp1* in the midbrain of 4 dpf *Tg(mrc1a:GFP)* wildtype (n=13) and *MZ*-/- (n=16) animals. **(N)** Quantification of the percent of *mrc1a*^+^ cells expressing *mpp1* in the midbrain of 4 dpf *Tg(mrc1a:GFP)* wildtype (n=13) and *MZ*-/- (n=17). **(O)** Quantification of the number of BAMs in 4 dpf *Tg(mrc1a:GFP)* animals injected with Cas9 or Cas9 + *mpp1* gRNA pool. **(P)** Agarose gel showing PCR on cDNA for *mpp1* and *tubulina1a* in 1 hpf WT and *MZ*-/- animals. **(Q)** Quantification of the number of *mrc1a*^+^ cells in the yolk sac of 1 dpf *Tg(mrc1a:GFP)* animals injected with Cas9 or Cas9 + *mpp1* gRNA pool. Scale bars equal 100 µm (C, H, J, and L). Error bars denote ± SEM (D, E, I, K, M, N, O, Q). GOI, gene of interest. The raw data for each group and statistical analyses are displayed in **supplemental table 2**.

### Validation of screen hits

To validate the hits from our high-throughput screen we performed additional gRNA injections and imaging. *Tg(mrc1a:GFP*) animals were injected at the one cell stage with Cas9 + gRNA pools corresponding to our 6 screen hits. Cas9-injected animals were used as controls. The number of *mrc1a*^+^; 4C4^+^ cells in the midbrain at 4 dpf was scored. Analysis was blinded and T7 endonuclease assay confirmed whether animals had indels near at least one gRNA site. Animals containing at least one indel were kept in the analysis. Scoring *mrc1a*^+^; 4C4^+^ microglia abundance and comparing to Cas9-injected animals revealed a significant reduction in *mrc1a*^+^; 4C4^+^ microglia abundance in *f11r.1* crispants (p<0.0001), *gas6* crispants (p=0.0023), and *mpp1* crispants (p=0.0281). (**Figure 4H-4I, Figure S4K**). These data suggest that *f11r.1, gas6,* and *mpp1* are required for proper *mrc1a*^+^ microglia abundance in the brain.

### *sox17* mutation alters transcription in *mrc1a*^+^ microglia

To better understand the genetic landscape of microglia development and how maternal *sox17* could be contributing to microglia abundance, we asked if the microglia phenotype observed in *sox17 MZ*-/- mutants was exacerbated by genetic perturbation of our 3 validated hits. We probed this question by creating crispants for our screen hit genes in the *MZ*-/- genetic background. To accomplish this, we injected Cas9 + a gRNA pool targeting *f11r.1, gas6,* or *mpp1* into wildtype and *MZ-/-Tg(mrc1a:GFP)* animals at the one cell stage. Cas9-injected animals of both genotypes and single mutant animals were used as controls. Analysis was blinded and T7 endonuclease assay was used to confirm which animals were crispants for the screen hit gene. Animals with an indel near at least one gRNA site were kept in the analysis. We scored the number of *mrc1a*^+^; 4C4^+^ microglia in the midbrain at 4 dpf and found that, aside from Cas9-injected controls (where *MZ*-/- mutants had significantly fewer *mrc1a*^+^; 4C4^+^ microglia in the brain, phenocopying earlier results), there were no differences in the number of *mrc1a*^+^; 4C4^+^ microglia in the brain between *f11r.1* crispants with or without an *MZ*-/- genetic background (p=0.9980) (**Figure 4J-4K**). We saw the same result for *gas6* and *mpp1* crispants with or without an *MZ*-/- genetic background (*gas6* p=0.5323, *mpp1* p=0.6054) (**Figure 4J-4K**). These results provide insight into the potential for undescribed gene regulatory networks governing microglia numbers during early stages of embryogenesis.

To ask about the potential cooperation of transcripts when *sox17* is perturbed, we investigated the co-expression of *sox17* transcript with *f11r.1, gas6,* or *mpp1* transcript in wildtype and *MZ-/-Tg(mrc1a:GFP)* animals using HCR on 4 dpf animals. Because *MZ*-/- mutants have fewer *mrc1a*^+^ cells, we scored the percent of *mrc1a*^+^ cells co-expressing *sox17* and *f11r.1, gas* or *mpp1*. Compared to wildtype animals in which we observed 35.00 ± 10.31% of *mrc1a*^+^ cells co-expressing *sox17* and *f11r.1, MZ*-/- mutants displayed 17.19 ± 7.93% of *mrc1a*^+^ cells co-expressing the two transcripts (p=0.2225) (**Figure S4L-S4M**). In assaying co-expression of *sox17* and *gas6* transcript, wildtype animals had 14.89 ± 10.92% of *mrc1a*^+^ cells co-expressing the two transcripts, compared to 22.83 ± 7.09% in *MZ*-/- animals (p=0.3670) (**Figure S4N-S4O**). These data show no change in transcript co-expression between wildtype and *MZ*-/- mutants. However, when analyzing co-expression of *sox17* and *mpp1* transcripts, we found that wildtype animals had 66.69 ± 11.79% of *mrc1a*^+^ cells co-expressing the two transcripts compared to only 15.25 ± 4.57% of *mrc1a*^+^ cells with co-expression in *MZ*-/- mutants (p=0.0002) (**Figure 4L-4M**).

Because s*ox17* is a transcription factor, we considered the hypothesis that *sox17* regulates *mpp1* mRNA levels since co-expression of the two transcripts were reduced in *MZ*-/- mutants. *mpp1* is membrane palmitoylated protein 1 that is a member of the MAGUK family of proteins^53^. These proteins interact with the cytoskeleton and can regulate cell proliferation, signaling pathways, and intercellular junctions^53^. We scored the percent of *mrc1a*^+^ cells expressing *mpp1* transcript in both wildtype and *MZ-/-Tg(mrc1a:GFP)* animals at 4 dpf using HCR. Compared to wildtype animals in which we observed 75.00 ± 11.32% of *mrc1a*^+^ cells expressing *mpp1, MZ*-/- mutants displayed a significantly lower percent (34.33 ± 6.75%) of *mrc1a*^+^ cells expressing *mpp1* transcript (p=0.0083) (**Figure 4L-4N**), supporting the idea that *sox17* positively regulates *mpp1* expression, and that this regulation is disrupted in animals lacking maternal contributions of *sox17*.

### *mpp1* perturbation does not impact BAM abundance

*sox17* mutants do not display differences in *mrc1a*^+^ BAM abundance at 4 dpf (**Figure 1H**), indicating that the maternal impact of *sox17* is specific to microglia. We next asked if genetic perturbation of *mpp1* impacts *mrc1a*^+^ BAM abundance at 4 dpf, with the hope of identifying genetic modifiers that may be involved in the specification of microglia vs BAMs. *Tg(mrc1a:GFP)* animals were injected with Cas9 + a gRNA pool targeting *mpp1* at the one cell stage, grown to 4 dpf, and stained with 4C4 to distinguish microglia from BAMs. Animals with an indel near an *mpp1* gRNA site were kept in the analysis. Cas9-injected animals were used as controls. Scoring the number of *mrc1a*^+^ cells at the brain borders revealed no difference in the number of BAMs between Cas9-injected animals (2.32 ± 0.33 BAMs) and *mpp1* crispants (2.41 ± 0.31 BAMs) (p=0.8436) (**Figure 4O**). Coupled with data from above, these results indicate that mutations in either *sox17* or *mpp1* do not impact the BAM population, but do specifically impact microglia.

### *mpp1* is maternally-contributed to offspring

Given that *sox17 MZ*-/- mutants show reduced *mpp1* transcription (**Figure 4N**), we asked if *mpp1* was also present before zygotic genome activation and could be contributing to the maternal effect. We tested if *mpp1* was maternally-contributed by collecting RNA from wildtype and *MZ-/-Tg(mrc1a:GFP)* animals at 1 hpf. Reverse transcription of RNA and amplification of the cDNA for a ∼250 bp region of the *mpp1* coding sequence showed a clear amplicon (**Figure 4P**, full gel image **Figure S4P**), indicating that *mpp1* is maternally-contributed from mother to offspring.

### *mpp1* regulates microglia abundance independently of yolk sac precursor abundance

Because *mpp1* phenocopies the reduction in microglia in *MZ*-/- mutants and is also maternally contributed (**Figure 4I, 4P**), we next asked if *mpp1* also regulated *mrc1a*^+^ yolk sac cells like *sox17*. Cas9 + the gRNA pool targeting *mpp1* was injected into the one cell stage of wildtype *Tg(mrc1a:GFP)* animals. Animals with an indel near an *mpp1* gRNA site were kept in the analysis and Cas9-injected animals were used as controls. Animals were grown to 1 dpf and the yolk sac was imaged. Scoring the number of *mrc1a*^+^ cells revealed 6.86 ± 0.60 *mrc1a*^+^ cells in the yolk sac of Cas9 controls compared to 5.94 ± 0.76 cells in *mpp1* crispants (p=0.3533) (**Figure S4Q**). We observed no difference in the number of *mrc1a*^+^ yolk sac cells in Cas9-injected controls vs *mpp1* crispants, supporting the idea that *mpp1* likely regulates microglia abundance independent of yolk sac precursors.

## DISCUSSION

The contribution of genetic material from mother to offspring is an essential component of proper embryogenesis^9^. Here, we identify *sox17* as a previously unrecognized maternal effect gene that is essential for microglia development in zebrafish. We show that *sox17* mRNA is maternally deposited and loss of maternal *sox17* leads to a reduction in *mrc1a*^+^ microglia throughout development, a phenotype potentially caused by a reduction of yolk sac precursors. Our CRISPR-based screen revealed 3 additional candidate genes that impact microglia, helping to identify that *mpp1* is expressed in *mrc1a*^+^ cells and co-expresses with *sox17* in *mrc1a*^+^ cells. Together, these findings reveal the role of *sox17* as a maternal effect gene with a postnatal phenotype in regulating microglia abundance, while also identifying *mpp1* as an additional genetic modifier of embryonic microglia abundance that is dependent on maternal *sox17* contribution.

### *sox17* as a maternal effect gene

There is some evidence that *sox17* can exhibit a maternal effect. In mice, maternal zygotic *sox17* mutants in which maternal *sox17* was reduced with oocyte-specific Cre recombinase (*Zp3::cre*) have a reduction in the overall inter cell mass (ICM) compared to zygotic mutants at E4.5^31^. Removal of maternal *Sox17* resulted in phenotypic changes in both homozygous and heterozygous embryos^31^, supporting an important contribution of maternal *Sox17* that is independent of the zygotic genotype. Additionally, the embryos of *Sox17* mutant mothers fail to implant properly into the uterus^54^. In our study, loss of maternal *sox17* caused a reduction in the number of *mrc1a*^+^; 4C4^+^ microglia in the brain. Importantly, this effect is phenocopied in maternal heterozygous (*MZ*+/-) animals, indicating that the absence of maternally-contributed transcript alone is sufficient to impact microglial abundance, regardless of the zygotic genotype.

We detected *sox17* transcript well before the maternal to zygotic transition. While maternal *sox17* loss reduced the number of *mrc1a*^+^ cells in the yolk sac at 1 dpf, rescue of *sox17* via injection of *sox17-P2A-mCherry-pA* RNA at the 1-cell stage restored both *mrc1a*^+^ cell numbers in the yolk sac and microglial abundance in the brain at 4 dpf in *MZ*-/- animals. These data directly demonstrate that mimicking maternal *sox17* transcript contribution is sufficient to restore microglia development, suggesting that *sox17* acts during an early, pre-zygotic genome activation window to regulate the abundance or survival of microglia. Interestingly, while microglia were significantly reduced in *MZ*-/- animals at both 4 and 14 dpf, *sox17* transcript expression in *mrc1a*^+^ cells was only reduced at 4 dpf and not 14 dpf. This finding suggests that while *sox17* transcript production eventually recovers to wildtype levels, early reduction is sufficient to cause long-term deficits in microglial populations. This distinguishes *sox17* from classical maternal effect genes, whose influence typically wanes after zygotic genome activation^50^, and instead positions it as a temporal bridge between maternal control and terminal cell fate in microglia development.

### Maternal effect genes impact terminally differentiated cells

Only a small number of maternal effect genes are known to be sublethal, including *auts2* and *fmr1* in fish, *gcl* in *Drosophila,* and *Kdm1a* in mice^3,11–13^. The *sox17* mutation reported here is likely hypomorphic, because null mutations in *sox17* result in embryonic lethality^23^. This brings up the interesting idea that hypomorphic alleles of essential genes may go unnoticed as potential maternal effect genes due to a lack of studies on their sublethal versions. There are specific examples that underscore this concept. Hypomorphic mutation of the essential gene *Cdc7*, which is critical for DNA replication, results in mice that are viable but exhibit growth defects and impaired gametogenesis^55^. Similarly, partial loss-of-function mutations in *Snrnp40*, a spliceosomal gene, cause adult mice to have severe immune dysfunction but no early lethality^56^. Such cases underscore the importance of studying sublethal alleles in the context of maternal effects to reveal late-onset or tissue-specific roles of these genes that were previously classified as strictly essential.

In humans, there are sublethal *SOX17* mutation variants that confer diseases of the lungs, heart, or kidneys^26–29^. Despite being a known regulator of vasculature, maternal mutants of *sox17* in this report did not disrupt midbrain vascular patterning, nor did it alter the number of BAMs^30,43^. BAMs share developmental progenitors with microglia, and both originate from early yolk sac hematopoiesis^17,18^. Transcriptional analysis and fate mapping studies in mice have begun to dissect the developmental program that distinguishes BAMs from microglia, although additional studies are needed to uncover the complex genetic landscape of their differentiation^18^. This report begins to unravel that genetic network by showing that only microglia, and not BAMs, are sensitive to *sox17* maternal loss and *mpp1* perturbation. This specificity may reflect differential transcriptional programs or epigenetic landscapes between microglia and BAM precursors that render microglia uniquely dependent on maternal *sox17* input.

### Maternal effect and microglia development

The idea that microglia are impacted by genetic maternal effect is novel and intriguing. This is, in part, because the impact of environmental maternal effects on microglia is well-documented^4^. Although microglia colonize the brain and become terminally differentiated in early development, the timing of this event is long after the maternal-to-zygotic transition of genome expression^34,50,52^. This means that a lack of maternally-contributed genetic material during early embryogenesis can have long-lasting effects in the embryo that impact critical cell populations, like microglia. Maternal RNAs begin to degrade at the onset of the maternal-to-zygotic transition of genome expression, and it is well documented that a majority of maternal transcripts in the embryo are degraded from 3.5-9 hpf^57,58^. Given these data, it is likely that little maternal transcript would remain by 4 dpf, the first time point that we observed reductions in microglia abundance. During maternal to zygotic transition, embryos rely heavily on maternal mRNAs to be translated into functional proteins that support essential processes^50^. Inhibition of translation during this stage disrupts the activation of zygotic transcription^50^. This mechanism may underlie our observation that *MZ*-/- mutants exhibit reduced *sox17* transcript levels at 4 dpf, well after zygotic genome activation and degradation of maternal mRNAs. These data suggest that maternal *sox17* contributes to the activation or maintenance of zygotic *sox17* expression. The role of maternal mRNAs in modulating the timing and robustness of zygotic gene expression highlights a broader regulatory network in which maternal transcripts help establish cell fate decisions even after zygotic genome activation.

Another structure well-positioned to be impacted by a maternal effect gene is the embryonic yolk sac. Microglia originate in the embryonic yolk sac^16,17^, a structure present at fertilization in zebrafish^51^, and therefore extremely likely to be influenced by maternal effect genes. Studies show that *Sox17* is expressed in fetal and neonatal hematopoietic stem cells and yolk sac hematopoiesis is reduced in *Sox17*-deficient mice^22^. Conditional knockout of *Sox17* showed a cell autonomous requirement for its expression in hematopoietic stem cells^22^. These experiments indicate that *Sox17* is required in hematopoietic stem cells for proper myeloid cell formation, providing yet another explanation for the impact of maternal *sox17* on microglia. Hematopoietic programs are also well known to be regulated by crosstalk between *Sox17* and the Wnt/B-catenin pathway^22^. Given that microgliogenesis is heavily reliant on the generation of hematopoietic stem cell microglia precursors^59^, these data align with the necessity for maternal contribution of *sox17* for embryonic yolk sac cell and microglia abundance. With the established role of *sox17* in yolk sac hematopoiesis^22^, and our finding of reduced *mrc1a+* yolk sac cells in *MZ*-/- mutants, we propose that maternal *sox17* activates zygotic *sox17* to promote the production of *mrc1a*^+^ yolk sac cells through Sox17/Wnt-regulated formation of hematopoietic stem cells. Coupled with our findings that *sox17* mutants have reduced *mpp1* transcription and *mpp1* crispants phenocopy the reduction in *mrc1a*^+^ microglia but not yolk sac cells, we hypothesize that *mrc1a*^+^ yolk sac cells colonize the brain and become microglia in a *mpp1-*dependent manner.

### Conclusion

Our discovery of *sox17* as a maternal-effect gene that governs terminally differentiated immune cell fates challenges existing paradigms about maternal contributions in vertebrates. It raises new questions about how other maternal-effect genes may influence immune cell development and how maternal environment or genetic regulation could affect immune ontogeny. Given the sensitivity of microglia to developmental perturbations and their central role in shaping brain wiring and function^15,41^, the maternal regulation of microglial number may have downstream consequences for neurodevelopmental processes and disorders. Understanding how maternal gene products interface with early hematopoiesis and microglia colonization of the brain may help explain susceptibility to disorders such as autism or schizophrenia, where microglial dysfunction has been implicated.

## LIMITATIONS

While our genetic interrogation revealed genes that impact the number of *mrc1a*^+^ microglia, it is not appropriate to conclude that the genes in the interrogation that did not generate a phenotype are not important for microglia abundance. Although our results reveal a reduction in yolk sac cells in *sox17* mutants that we presume contributes to the reduction in microglia, tracking these cells from the yolk sac to the brain to uncover how *sox17* regulates *mrc1a*^+^ yolk sac cells and microglia number is beyond the scope of this study. Our results indicate that sublethal *sox17* mutants have reduced *mpp1* transcript levels, but the data does not distinguish between direct vs indirect control. We could not identify consensus *sox17* binding regions in the *mpp1* promoter region, but enhancer regions might contain such binding sites. Alternatively, *sox17* could indirectly regulate *mpp1* through an additional transcriptional target. We hypothesize that the protein generated from the sublethal *sox17* mutation likely retains some functionality because it is not lethal, but we do not know the specific impacts on protein function.

## ACKNOWLEDGEMENTS

We thank F. Lin (University of Iowa) for sending us the *Tg(sox17:H2A-mCherry)* animals. We thank current and previous members of the Smith lab for their insightful discussions. We especially thank Dana DeSantis who managed and established the screening pipeline in the lab. We owe gratitude to Rebecca Wingert who brought our attention to the uniqueness of the maternal effect phenotype on microglia. We would also like to thank 3i for answering any imaging-related questions and Deborah Bang and other animal facility staff for zebrafish housing and upkeep. This paper was supported by the University of Notre Dame, the Elizabeth and Michael Gallagher Family, Centers for Zebrafish Research and Stem Cells and Regenerative Medicine at the University of Notre Dame, the SMART foundation (CJS), Chan Zuckerberg Initiative (CJS), and the NIH (CJS: DP2NS117177, CAH: F31HD111206, F99NS139552).

## METHODS

### Experimental model and subject details

All experimental procedures adhered to the NIH guide for the care and use of laboratory animals and was approved by The University of Notre Dame Institutional Animal Care and Use Committee (Protocol #22-07-7322). Notre Dame IACUC adheres to the United States Department of Agriculture, the Animal Welfare Act (USA), and the Assessment and Accreditation of Laboratory Animal Care International.

Zebrafish stable strains used in this study were *Tg(mrc1a:GFP)*^60^, *Tg(mrc1a:GFP); sox17^nt206^,* (details of generation below), *Tg(pu1:Gal4;UAS:GFP)*^61^, and *Tg(sox17:H2A-mCherry)*^62^. Zebrafish embryos were produced through pairwise mating and placed in an incubator at 28°C in constant darkness. Animals were staged by hpf and dpf^63^. Beginning at 24 hpf, embryos were treated with PTU (0.003%) to reduce pigmentation for imaging. For experiments in which 14 dpf animals were used, embryos were screened for GFP and sent back to the zebrafish facility at 5 dpf and maintained by our veterinarian technicians until 14 dpf. At 14 dpf, immunohistochemistry or hybridized chain reaction was performed as stated.

### CRISPR/Cas9 knockdown

Guide RNAs (Integrated DNA Technologies) against genes of interest were generated using CHOPCHOP and injected into *Tg(mrc1a:GFP)* animals at the one cell stage. For single guide injections (*sox17* crispants), the injection mix also contained 5 µM AltR S.p. Cas9 Nuclease V3 (Integrated DNA Technologies), as previously done^64^. For pooled gRNA injections (high throughput screen), the injection mix also contained 2 µM EnGen Spy Cas9 NLS (New England Biolabs).

*Sox17* gRNA sequences

Exon 1: 5’ GTCACTGGAGTACCCCGCAT 3’

Exon 2: 5’ ACCCCGGCAAAGGTGCACTG 3’

### T7 endonuclease assay

To determine if animals had an indel in or near a gRNA site a T7 endonuclease assay was performed (New England Biolabs) as previously done^65,66^. The assay was carried out according to the manufacturer’s instructions. In brief, genomic DNA was individually extracted from animals injected with gRNA and a small region around the gRNA target site was amplified by PCR. This PCR produced was briefly annealed in a reaction containing NEB Buffer 2 and then T7 Endonuclease was added to the reaction. Digested products and original PCR products were run side-by-side using gel electrophoresis. Only animals with digested PCR product (indicating an indel) were kept in the analysis.

### Generation of the 4C4 antibody

The anti-4C4 antibody was created by collecting supernatant of 7.4.C4 mouse hybridoma cell cultures. Cells were obtained from the European Collection of Authenticated Cell Cultures (92092321) and grown in accordance with their suggested protocol. Briefly, cells were grown in suspension at 37°C in 8% CO2 in a culture media containing DMEM + 2mM glutamine + 1% (v/v) Hypoxanthine, Thymidine (HT) supplement (100x concentrate containing 10mM sodium hypoxanthine and 1.6mM thymidine) + 10% Fetal Bovine Serum (FBS). Cells were supplemented with 20% fetal bovine serum (FBS) until actively proliferating cultures were obtained, and then the FBS concentration was reduced to 10% of the media volume. Cells were pelleted for 5 min at 100 x g and the supernatant was collected and aliquoted for use in immunohistochemistry.

### Immunohistochemistry

The same immunohistochemistry protocol (adapted from the Hibi lab) was performed on all tissue samples as done previously^64,65^. The primary antibody used was anti-4C4 (1:50). The secondary antibody used was Alexa Fluor 647 goat anti-mouse IgG H+L (1:600 Thermo Fisher). Larvae were fixed using 4% PFA in PBS with 0.1% Triton X-100 at 25°C for 1-2 hours. Larvae were then washed with PBST (PBS, 1% TritonX-100) for 5 minutes followed by a 5 minute wash with DWTx (distilled water, 1% Triton). Larvae were then incubated with cold acetone and chilled at −20°C for 10 minutes and washed with PBST 3 × 5 minutes. Next, larvae were incubated for an hour in 5% goat serum in PBST at 25°C. Larvae were then incubated in 5% goat serum in PBST with the primary antibody for an hour at 25°C and then overnight at 4°C. After 3 washes of PBST for 30 minutes each, the larvae were then incubated in 5% goat serum in PBST with the secondary antibody at 25°C for an hour and then transferred to 4°C overnight. The larvae were then washed 3 times with PBST for 30 minutes and finally transferred to a 50% glycerol/PBST solution and stored in 4°C until imaging was performed.

### *In vivo* imaging

For imaging, animals were anesthetized with 1X tricaine-S (Pentair Aquatic Ecosystems) and mounted dorsally (midbrain imaging) or laterally (yolk sac imaging) in 35 mm glass-bottom dishes with 0.8% low melting point agarose. One the agarose was solidified, dishes were filled with 1X tricaine-S. Images were acquired using a custom built Intelligent Imaging Innovations (3i) spinning disc confocal microscope with a Zeiss Axio Observer Z1 Advanced Mariana Microscope, X-Cite 120LED White Light LED System, filter cubes for GFP and mCherry, a motorized X,Y stage, a Piezo Z stage, 20× Air (0.50 NA), 63× (1.15 NA), 40× (1.1 NA) objectives, CSU-W1 T2 Spinning Disk Confocal Head (50 μM), Teledyne Prime 95B sCMOS camera, dichroic mirrors for 446, 515, 561, 405, 488, 561, and 640 excitation, laser stack with 488 nm, 561 nm, and 637 nm with laser stack FiberSwitcher, and Ablate Photoablation System (532 nm pulsed laser; pulse energy 60 J at 200 Hz) as done previously^64–66^. All images were acquired using the most current version of 3i’s Slidebook software to capture a 100 μm z stack range with a 1μm step size. The z stack was acquired by setting a position at the dorsal most point of the midbrain or yolk sac and imaging 100 μm from that point. Images were processed using Slidebook and ImageJ. Only brightness and contrast were adjusted and enhanced for images represented in this study.

### Microglia and BAM quantification

*mrc1a*^+^, 4C4^+^, and *mrc1a*^+^; 4C4^+^ cells in the brain were quantified using Slidebook’s 3D volume viewer to count the number of cells present in the midbrain region. Because perivascular cells that label with *mrc1a* also exist in the midbrain, we only included cells for analysis that were freely floating and not in contact with *mrc1a*-labeled vasculature as done previously^65^. For BAM quantification, the number of *mrc1a*^+^ cells outside the brain borders were counted.

### Generation of *sox17^nt206^* stable mutant line

*Tg(mrc1a:GFP)* animals were injected (F0 generation) at the one cell stage with a gRNA targeting exon 2 of the gene. Animals were grown to adulthood and outcrossed in a pairwise manner to AB fish. 20 embryos (F1 generation) per clutch were collected and lysed for DNA collection. Lysed DNA was pooled and a 250 bp region containing the gRNA target site was sequences. Of the F0 animals, the ones that gave rise to offspring showing non-wildtype sequence in the region of interest were outcrossed again to AB animals and F1 animals were grown to adulthood. Once adults, DNA was collected from F1 animals for sequencing. Sanger sequencing revealed an animal with an eight-base pair deletion in exon 2 of *sox17* resulting in an early stop codon and predicted protein truncation. This deletion disrupted the recognition site for ApaLI so animals were able to be genotyped using restriction digest. The F1 animal with the eight-base pair deletion was outcrossed to AB, embryos were grown to adulthood, genotyped, and tanks of heterozygous animals were maintained. Homozygous tanks were also generated by incrossing heterozygous animals.

### Vasculature integrity

To score the integrity of the vasculature in and around the midbrain, ImageJ was used to quantify length, area, and area covered by signal. For the length of the midcerebral vein, a line was drawn laterally through the visible fluorescence of the midcerebral vein^45^ (**figure S1E, label F**) and the length of that line was measured in ImageJ. For the length of the mesencephalic vessels^45^, a line was drawn from the point in the maximum z-projection where the mesencephalic vessels branch on the dorsal side of the head to extend laterally towards the vascular loops (**figure S1E, yellow G**). The length of each line was measured in ImageJ and the sum of these two lines is plotted. To measure the sum of the area of the vascular loops^46^, the polygon tool in ImageJ was used to trace the cells comprising the vascular loops that sit dorsolateral to the midbrain (**figure S1E, yellow H**). The sum of the area of these loops was measured in ImageJ and plotted. We used vascular GFP^+^ signal as a proxy for the overall amount of vasculature present in each image. To measure this, we used the ImageJ area fraction analysis tool with a threshold of 219-655355. These thresholds were chosen because they delineated between images where fish had little to no autofluorescence in the skin versus images where fish had excessive autofluorescence in the skin (**figure S1E**). ImageJ calculated the percent of the image covered by signal. Animals that fell outside of the threshold were excluded from analysis.

### Whole mount Hybridized Chain Reaction

HCR in whole mount zebrafish embryos was performed using the Molecular Instruments Inc. protocol reagents and probes as done previously^64^. Briefly, animals were fixed with 4% paraformaldehyde in PBST for one hour. Animals were then dehydrated with 100% methanol (MeOH) and stored at -20°C overnight. The next day, animals were rehydrated with a series of MeOH and PBS with 0.1 % Tween 20 (PBSTw) washes for five minutes each (75% MeOH/25% PBSTw, 50% MeOH/50% PBSTw, 25% MeOH/75% PBSTw, and 100% PBSTw). Animals were permeabilized with 30 µg/mL of proteinase K at room temperature for 35 minutes (4 dpf) or 45 minutes (14 dpf) followed by two room temperature washes with PBSTw without incubation. Animals were permeabilized with 30 µg/mL of proteinase K at room temperature for 35 min followed by two washes at room temperature with PBSTw without incubation. A postfix was performed by placing animals in 4% paraformaldehyde in PBST for 20 minutes at room temperature followed by 5 × 5 min washes with PBSTw. Hybridization of the probe corresponding to the mRNA sequence of interest was done by incubating animals with the probe (1:250) at 37 °C overnight. The next day, excess probe was washed away and animals were incubated overnight at room temperature with the amplifier (30 pmol, B1 or B3). The following day, excess amplifier solution was washed off with 5x sodium chloride sodium citrate and animals were stored in PBSTw at 4 °C until imaging. HCR fish were protected from light during the HCR protocol and storage prior to imaging.

### Maternal contribution of genetic material

To confirm if mRNA was maternally contributed to offspring, we collected RNA from embryos at 1 hpf using the Zymo Direct-zol RNA Miniprep kit (cat#R2051). RNA was reverse transcribed using the SuperScript IV kit (Invitrogen) to obtain cDNA. PCR amplicons for each gene were generated using cDNA as starting material and *tubulina1a* was used as the loading control. PCR products were run on a 1% agarose gel at 120V for 45 minutes.

*sox17* primer sequences

F 5’ TGAACTGCCCATCAACTTTAGA 3’

R 5’ TGCTCAAGAATTATTATAGCCGC 3’

*mpp1* primer sequences

F 5’ AACAAGAAGGATCCCAACTGGTGG 3’

R 5’ TCTGTTGAACGCTGGAAGTCTCAC 3’

### *sox17* rescue

The *sox17* coding sequence was reverse transcribed from 1 dpf cDNA. We used HiFi to insert the *sox17* coding sequence into a *p3e-Cas9-p2A-mCherry-pA* construct, from which the Cas9 had been removed, to create a *p3e-sox17-p2A-mCherry-pA* construct. This construct was linearized and *in vitro* transcribed using the mMESSAGE mMACHINE T7 kit (Invitrogen). RNA was injected at the one cell stage and embryos were screened for mCherry expression at 24 hpf to confirm translation of *sox17*.

### Single cell sequencing

The microglia scRNAseq data were downloaded from GSE121654^32^. Data pre-processing, normalization, and clustering was performed using the R package Seurat (v. 5.3.0)^67^. Single-cell transcriptomes were initially filtered for quality control. Cells with fewer than 200 genes detected and genes expressed in fewer than three cells were removed. Data normalization was performed using the function NormalizeData with normalization.method = “LogNormalize” and scale.factor = 10,000. 2,000 variable genes were chosen using the function FindVariableFeatures with selection.method = “vst.” Data scaling was performed using the ScaleData function. Dimensional reduction was accomplished by performing principal component analysis and then using the first 20 principal components for Uniform Manifold Approximation and Projection (UMAP) using default parameters associated with the RunUMAP function. Unsupervised clustering was finished by constructing a shared nearest neighbor (SNN) graph using the FindNeighbors function and then performing graph-based clustering using the “Louvain” algorithm with resolution = 0.5, resulting in 14 clusters. Differential expression analysis between clusters was performed using a Wilcoxon rank sum test by the FindAllMarkers function. The violin plots and feature plots for chosen marker genes were obtained by the VlnPlot and FeaturePlot functions, respectively.

### High throughput CRISPR/Cas9 genetic screen

For each of the 26 genes chosen from single cell sequencing analysis for screening, 4-6 gRNAs were generated using CHOPCHOP. For unduplicated genes 4 gRNAs were generated, while 6 gRNAs were generated for duplicated genes (3 gRNAs/gene duplicate). Selection followed these parameters: guides should ideally be 20bp in length, begin with GA, and have no off-targets with fewer than 3 bp mismatches. A list of genes and their guides are included in **supplemental Table 1.**

*Tg(mrc1a:GFP)* animals were injected at the one cell stage with an injection mix containing the gRNA pool, 2 µM EnGen Spy Cas9 NLS (New England Biolabs), phenol red, and nuclease-free water. Injected animals were screened for GFP at 4 dpf and placed in a 96-well plate with 1X tricaine-S. Images were captured on a custom-built VAST robot coupled with a Zeiss Examiner.Z1 spinning disc confocal from 3i. A minimum of 7 animals were imaged and analyzed for each gene in the initial screen. For validation of the screen hit genes, T7 endonuclease assay was performed to confirm crispants and a minimum of 10 animals were included in the analysis.

### Generation of crispants in an *MZ*-/- genetic background

To generate double mutants *Tg(mrc1a:GFP); sox17nt206*-/- animals were incrossed to create maternal zygotic homozygous mutants. These embryos were injected at the one cell stage with Cas9 or Cas9 + gRNAs corresponding to our 3 validated screen hits. Animals were grown to 4 dpf and imaged as described. After imaging, animals were unmounted into individual strip tubes and T7 endonuclease assay was performed. Only animals with indels were kept in the analysis.

### Blind quantifications and statistical analysis

Scientists were blind to the identity of the experimental versus control group during quantifications whenever possible. Normality distribution of experiments was assessed using the “Normality and Lognormality Test” feature on Prism. When applicable, unpaired t-tests were performed. When the data did not meet the assumption for a normal Gaussian distribution, a Mann-Whitney test was performed. When applicable, a one-way ANOVA was performed, followed by Tukey’s post hoc test to account for variance. When the data did not meet the assumption for a normal Gaussian distribution, a Kruskal-Wallis test was performed followed by Dunn’s multiple comparisons test. When applicable, a 2way ANOVA with Šídák’s multiple comparisons test was performed. Descriptive stats were run on each set of data and the mean ± SEM was plotted.

### Software

Adobe Illustrator, Slidebook, GraphPad Prism, ImageJ, Google Colab and posit Cloud were used in this study to analyze, acquire, and compile figures. ChatGPT and Gemini aided in reviewing parts of the writing for conciseness and literature search.

### Contributions

C.A.H. performed the experimentation, analysis, writing, and editing of the manuscript. J.D. performed experimentation for the high throughput genetic screen. D.G. and J.L performed all single cell sequencing analysis and generation of UMAP and violin plots. C.J.S wrote and edited the manuscript and supervised and funded the project.

## Code availability

All code for the scRNA-seq analysis can be accessed at https://github.com/RavenGan/Microglia_analysis

## Contact for Reagent and Resource Sharing

Data that supports the findings of this study is available from the authors upon request. All data collected for the study is included in the figures.

---

**Supplementary Figure 1.**
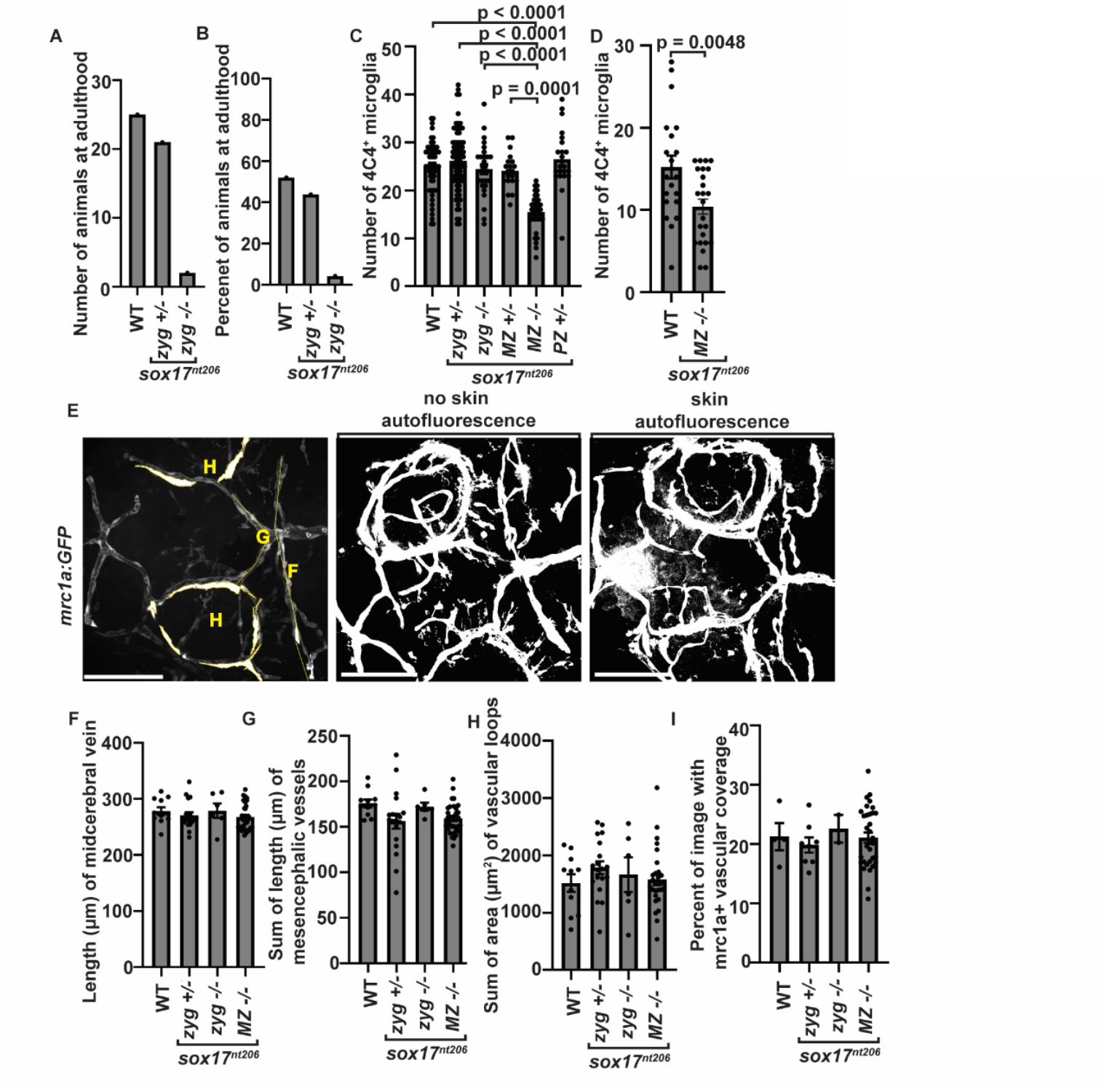
**(A)**Quantification ofthe number of animals corresponding to each genotype that survived to adulthood, generated from a F1 incross of *zyg*+/- mutants. **(B)** Quantification of the percent of animals at adulthood corresponding to each genotype generated from a F1 incross of *zyg*+/- mutants. **(C)** Quantification of the number of 4C4^+^ microglia in the midbrains of 4 dpf *Tg(mrc1a:GFP)* wildtype (n=55), *zyg*+/-(n=76), *zyg*-/- (n=28), *MZ*+/- (n=20), *MZ*-/- (n=37), and *PZ*+/- (n=23) animals. **(D)** Quantification of the number of 4C4^+^ microglia in the midbrains of 14 dpf *Tg(mrc1a:GFP)* wildtype (n=21) and *MZ*-/- (n=24) animals. **(E)** Confocal z-projections of 4 dpf *Tg(mrc1a:GFP)* wildtype animals showing the regions measured in F, G, and H, as well as an animal with vs without skin autofluorescence for animals measured in I. **(F)** Quantification of the length of the midcerebral vein in 4 dpf *Tg(mrc1a:GFP)* wildtype (n=11), *zyg*+/- (n=19), *zyg*-/- (n=6), and *MZ*-/- (n=33) animals. **(G)** Quantification of the sum of the length of the mesencephalic vessels in 4 dpf *Tg(mrc1a:GFP)* wildtype (n=11), *zyg*+/- (n=19), *zyg*-/- (n=6), and *MZ*-/- (n=33) animals. **(H)** Quantification of the sum of the area of the vascular loops in 4 dpf *Tg(mrc1a:GFP)* wildtype (n=11), *zyg*+/- (n=19), *zyg*-/- (n=6), and *MZ*-/- (n=28) animals. **(I)** Quantification of the percent of the image covered by GFP^+^ signal in 4 dpf *Tg(mrc1a:GFP)* wildtype (n=11), *zyg*+/- (n=19), *zyg*-/- (n=6), and *MZ*-/- (n=33) animals. Scale bars equal 100 µm (E). Error bars denote ± SEM (C, D, F, G, H, and I). The raw data for each group and statistical analyses are displayed in **supplemental table 2**.

**Supplementary Figure 2.**
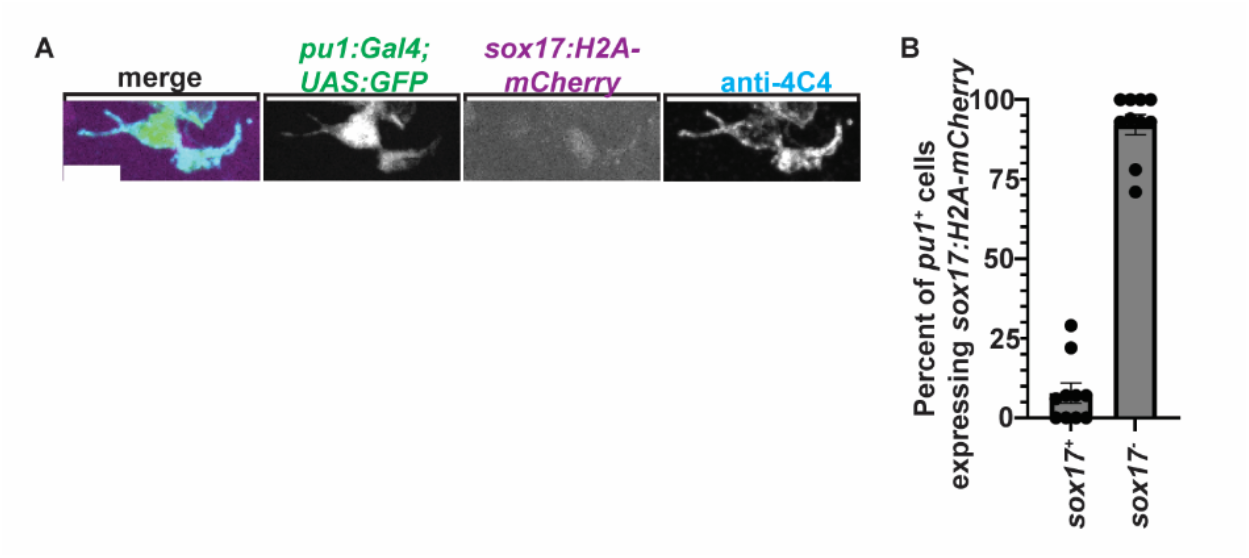
**(A)** Confocal z-projections of a single cell in the midbrain of a 4 dpf wildtype *Tg(pu1:Gal4; UAS:GFP); Tg(sox17:H2A-mCherry)* animal stained with 4C4. **(B)** Quantification of the percent of *pu1*^+^ cells that are *sox17:H2A-mCherry*^+^ and *sox17:H2A-mCherry^-^* in the midbrain of 4 dpf wildtype *Tg(pu1:Gal4; UAS:GFP); Tg(sox17:H2A-mCherry)* animals (n=10). Scale bar equals 10 µm (A). Error bars denote ± SEM (B). The raw data for each group and statistical analyses are displayed in **supplemental table 2**.

**Supplementary Figure 3.**
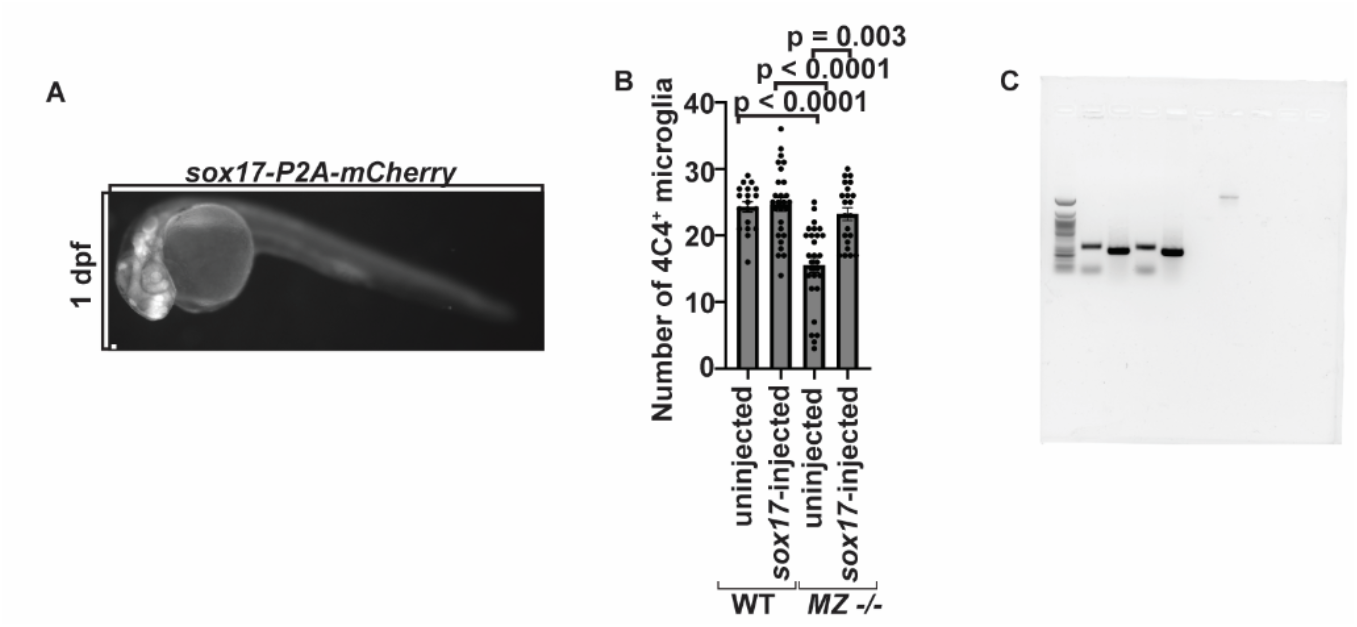
Zeiss Axiozoom image of a 1 dpf *Tg(mrc1a:GFP) MZ*-/- animal injected with *sox17-P2A-mCherry-pA* RNA at the one cell stage. **(B)** Quantification of the number of 4C4^+^ microglia in the midbrain of 4 dpf *Tg(mrc1a:GFP)* uninjected WT (n=19), injected WT(n=27), uninjected *MZ*-/- (n=28), and injected *MZ*-/- animals (n=22). **(C)** Uncropped image of agarose gel in **Figure 3A** showing PCR on cDNA for *sox17* and *tubulina1a* in 1 hpf WT and *MZ*-/- animals. Scale bar equals 100 µm (A). Error bars denote ± SEM (B). The raw data for each group and statistical analyses are displayed in **supplemental table 2**.

**Supplementary Figure 4.**
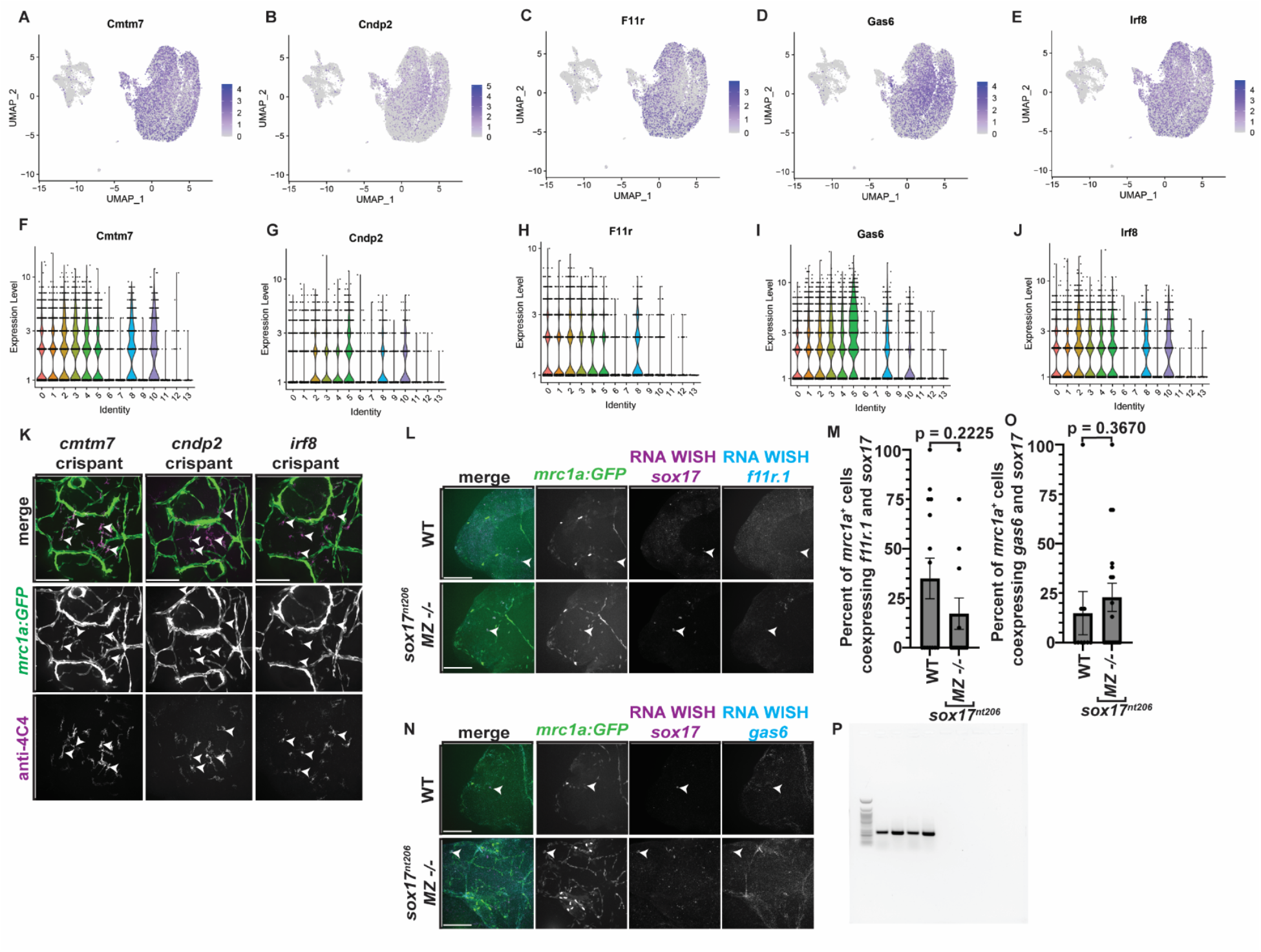
**(A)** UMAP of **(B)** *Cmtm7*, **(C)** *Cndp2,* **(D)** *Frr1,* and **(E)** *Irf8* expression across all 13 clusters^32^. Violin plot of **(F)** *Cmtm7*, **(G)** *Cndp2,* **(H)** *Frr1,* and **(I)** *Irf8* expression across all 13 clusters^32^. **(K)** Confocal z-projections of the midbrains of 4 dpf *Tg(mrc1a:GFP)* uninjected, Cas9-injected, Cas9 + *cmtm7* gRNA pool-, Cas9 + *cndp2* gRNA pool-, and Cas9 + *irf8* gRNA pool-injected animals stained with 4C4. White arrowheads point to *mrc1a*^+^; 4C4^+^ microglia. **(L)** Confocal z-projections of the midbrains of 4 dpf *Tg(mrc1a:GFP)* animals stained with WISH HCR probes against *sox17* and *f11r.1*. White arrowheads point to *mrc1a^+^; sox17^+^; f11r.1*^+^ cells. **(M)** Quantification of the percent of *mrc1a*^+^ cells co-expressing *sox17* and *f11r.1* in the midbrain of 4 dpf *Tg(mrc1a:GFP)* wildtype (n=14) and *MZ*-/- (n=16) animals. **(N)** Confocal z-projections of the midbrains of 4 dpf *Tg(mrc1a:GFP)* animals stained with WISH HCR probes against *sox17* and *gas6*. White arrowheads point to *mrc1a^+^; sox17^+^; gas6*^+^ cells. **(O)** Quantification of the percent of *mrc1a*^+^ cells co-expressing *sox17* and *gas6* in the midbrain of 4 dpf *Tg(mrc1a:GFP)* wildtype (n=9) and *MZ*-/- (n=18) animals. **(P)** Uncropped image of agarose gel showing PCR on cDNA for *mpp1* and *tubulina1a* in 1 hpf WT and *MZ*-/- animals. Scale bars equal 100 µm (K, L, and N). Error bars denote ± SEM (M and O). The raw data for each group and statistical analyses are displayed in **supplemental table 2**.

## Notes

### Competing Interest Statement

The authors have declared no competing interest.

